# Peripheral inflammation mediates midbrain Lrrk2 kinase activity via Rab32 expression

**DOI:** 10.1101/2025.09.05.674552

**Authors:** Jordan Follett, Isaac Bul Deng, Robert C. Sharp, Shannon Wall, Adamantios Mamais, Matthew J. Farrer

## Abstract

Mutations that increase leucine-rich repeat kinase 2 (LRRK2) activity confer significant risk for Parkinson’s disease (PD), yet incomplete disease penetrance suggest additional factors are required to manifest disease. We recently identified RAB32 Ser71Arg as a Mendelian gene for PD. Here, we establish Rab32 as a key mediator linking peripheral inflammation to Lrrk2 activation. We show that Rab32 and Rab38 expression are modestly, but inversely, correlated with their homolog Rab29. *In vivo*, peripheral lipopolysaccharide (LPS)-induced inflammation selectively induced Rab32 expression in midbrain Iba1^+^ microglia but not dopaminergic neurons, where it localized to Lamp1^+^ lysosomal compartments and correlated with Lrrk2 kinase activity. LPS similarly induces Rab32 expression in human induced pluripotent stem cell-derived microglia, demonstrating a unified biological response to inflammation across species. Promoter analysis identified Tfe3, a master regulator of lysosomal biogenesis and autophagy, as a key driver of Rab32 expression induced Lrrk2 kinase activation. During inflammation, Tfe3 translocated to the nucleus of midbrain Iba1^+^ microglia to induce Rab32 expression and Lrrk2 kinase activity. Knockdown of Tfe3, but not Tfeb, mitigates these effects, establishing Rab32 as a physiological rheostat of Lrrk2 activity. This mechanistic pathway enables peripheral inflammation to modulate LRRK2 activity and highlights Rab32/Tfe3 as a therapeutic targeting for neuroprotection in PD.

## INTRODUCTION

Parkinson disease (PD) is a multifactorial, neurodegenerative movement disorder, associated with the loss of striatal dopaminergic (DA) neurons emanating from the *substantia nigra pars compacta* (*SNpc*), with gliosis and alpha-synucleinopathy (*1*). Mutations in leucine-rich repeat kinase 2 *(LRRK2)* lead to a clinical syndrome that is indistinguishable from idiopathic PD. LRRK2 Gly2019Ser is most frequent and accounts for 13 to 30% of PD in Ashkenazi Jews and North African Berbers, respectively (*2*). LRRK2 PD is inherited ‘identical-by-descent’ from few ancestral founders (*3*) although the disease often manifests sporadically given its reduced age-associated penetrance which is population-specific (*2*).

All pathogenic LRRK2 substitutions that cause PD increase its kinase activity yet monomeric LRRK2 is largely inactive in the cytosol. Activation requires membrane association and dimerization via homologous Group II RAB GTPases, namely RAB29 (formerly RAB7L1), RAB32 and RAB38. All three interact with LRRK2 via its N’ terminal ankyrin domain (*4*) in which variability is inversely associated with disease risk (*5*). *In vitro*, Rab29 overexpression readily induces LRRK2 kinase activity (*6*) that has the propensity to phosphorylate 10-14 Rab GTPase substrates that dysregulates their localization and function (*7*). Polymorphic variants in *RAB29* are weakly associated with PD and correlate with its expression (*8*). Nevertheless, endogenous Rab29 does not impact basal nor stimulate Lrrk2 kinase activity *in vivo* (*9*). Thus, the physiologic factors responsible for LRRK2 kinase activity are incompletely defined.

Recently, RAB32 Ser71Arg was linked and associated with PD, and shown to increase LRRK2 kinase activity. Despite its global diaspora, the mutation is also inherited ‘identical-by-descent’ from one ancestral founder (*10*). RAB32 is acutely responsive in pathogen defense and immune signalling (*11*), and otherwise coordinates mitochondrial, lysosomal and metabolic activity to maintain cellular homeostasis (*12, 13*). Together, these processes converge on the Microphthalmia/Transcription Factor E (MiT/TFE) family, comprising TFEB, TFE3, MITF, and TFEC, which are dephosphorylated and translocate to the nucleus to control gene expression within the CLEAR (Coordinated Lysosomal Expression and Regulation) network. Its consensus sequence is a palindromic 10-bp GTCACGTGAC motif arranged in tandem within ∼200bp of regulated transcriptional start sites (*14*). TFEB and TFE3 are best described to initiate lysosomal biogenesis in response to inflammation to reinforce antimicrobial and degradative capacity. LRRK2 was recently proposed to act as a signaling hub to coordinate membrane trafficking with MiT/TFE localization (*15*) but whether RAB29, RAB32 and RAB38 affect LRRK2 activity and MiT/TFE distribution remain unexplored.

In this study, we assess Rab29, Rab32 and Rab38 gene and protein expression in mouse brain, including the relationship between these homologs with Lrrk2 kinase activity under basal conditions and with peripheral immune stimulation. We compliment this biology with human induced pluripotent stem cell-derived microglia. In a body-to-brain model of inflammation, we detail Rab32 as an immune-responsive GTPase that stimulates Lrrk2 kinase activity.

Furthermore, we demonstrate Tfe3 regulates Rab32 expression and thereby modulates Lrrk2 kinase activation. This establishes a mechanistic framework to target Tfe3 and thereby attenuate pathogenic Lrrk2 signaling.

## RESULTS

### Rab29 knockdown promotes LRRK2 activity via Rab32 expression

We evaluated commercial antibodies raised against Rab32, Rab38 and Rab29 in a murine cell line, given the close sequence homology of Group II sub-family of GTPases. While the siRNA-mediated loss of Rab32, Rab38 or Rab29 protein confirmed their specificity, there was a significant, inverse relationship between the levels of Rab29 and those of Rab32 and Rab38 **(Fig.1).** Knock down of Rab32 protein levels to <0.01 of that observed in NTC cells did not alter Rab29 (p=0.81) or Rab38 (p=0.99) (compare **Fig.1Ba-c**). Similar knock down of Rab38 protein to <0.10 of that observed in NTC cells was accompanied by a 0.3-fold (±S.E.M 0.10) non-significant decrease in Rab32 (p=0.10), although Rab29 levels remained unchanged (p=0.18; compare **Fig.1Bc** with **Ba** and **Bb**). However, knock down of Rab29 protein to <0.05 of that observed in NTC cells lead to significant 1.6-fold (±S.E.M 0.07) increase in Rab32 (p=0.0003) and 1.25-fold (±S.E.M 0.06) increase in Rab38 (p=0.0002; compare **Fig.1Bb** with **Ba** and **Bc).**

**Fig. 1.**
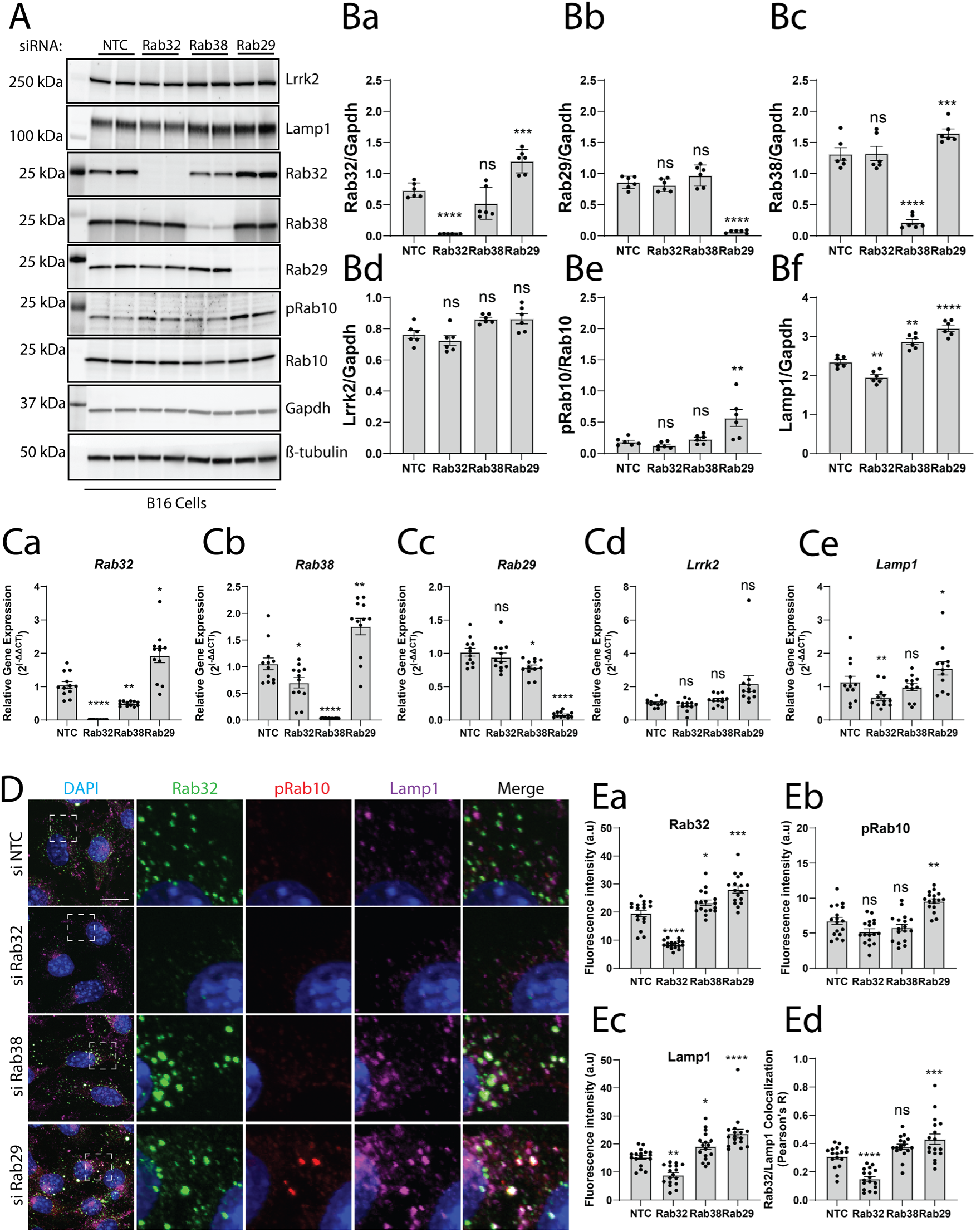
Rab29 knockdown promotes LRRK2 activity via Rab32 expression. **A)** Representative western blot data of siRNA-mediated antibody validation in B16 cells for Rab32-sub family of GTPases, Lrrk2, Lamp1 and pT73-Rab10/Rab10. Data is representative of three independent experiments. **B)** Quantification of protein lysates analyzed in (**A)**. **C)** Relative gene expression (2^(-ΔΔCT)^) of *Rab32*-sub family of GTPases in B16 cells following siRNA-mediated knock down. Data is pooled from three independent experiments where each dot on the graph represents an individual well that underwent the treatment paradigm. **D)** Representative confocal microscopy of Rab32, pT73-Rab10 and Lamp1 in B16 cells following siRNA-mediated knock down. Nuclei are counterstained with DAPI. Images are from three independent experiments with 4-7 images captured per experiment. Scale bar = 10µm. **E)** Quantification of fluorescence intensity and co-localization (Pearson’s R) analyzed in (**D).** All data are presented as mean ± S.E.M. Statistical analysis was determined through a one-way ANOVA with a Tukey multiple comparisons *post-hoc* test, where significant findings are reported relative to NTC. *\*p* <0.05, *\*\*p*<0.01, *\*\*\*p*<0.001, \**\*\*\*p*<0.0001.

There was no significant difference in total Lrrk2 with any of these siRNA treatments (**Fig.1Bd & Cd;** p = 0.70, 0.06 & 0.06, for siRNA *Rab32*, *38* and *29*, respectively). The phosphorylation of Rab10 at a known Lrrk2 phosphorylation site (pThr73-Rab10) remained comparable to NTC levels with knock down of Rab32 (p=0.93) and Rab38 (p=0.97) but was 2-fold increased (±S.E.M 0.13; p=0.005) with the knock down of Rab29 (**Fig.1Be**). While this may seem contrary to the literature (*9*), the activation of Lrrk2 kinase observed is consistent with the modest, reciprocal elevation in Rab32 and Rab38 when Rab29 is knocked down (**Fig.1Ba and 1Bc**).

Lamp1 levels were significantly diminished 0.18-fold (±S.E.M 0.06) compared to NTC cells with Rab32 knock down (p=0.005), whereas Lamp1 levels significantly increased 1.25 (±S.E.M 0.11) and 1.34-fold (±SEM 0.13) with Rab38 (p=0.005) and Rab29 (p<0.0001) knock down, respectively (**Fig.1Bf**).

Complementary observations were made by qPCR that confirmed siRNA-mediated loss of *Rab32*, *Rab38* and *Rab29*, albeit with no significant impact on the expression of *Lrrk2* (**Fig.1C**; p=0.55, 0.43 & 0.13 for siRab32, 38 and 29, respectively). Modest differences in gene expression were observed for individual Rabs that reflect observations made for their respective proteins; *Rab29* knock down leads to a significant increase in *Rab32* (p=0.01) and *Rab38* (p=0.006) gene expression, suggesting transcriptional control (compare **Fig.1Ca** and **Cb**). The knock down of *Rab38* modestly decreased the expression of *Rab32* (p =0.001) and *Rab29* (p=0.01) **(Fig.1Ca** and **Cc**), while the knock down of *Rab32* decreased *Rab38* (p=0.02) but not *Rab29* (p=0.72) expression (**Fig.1Cb** and **Cc**). Lastly, the knock down of *Rab32* (p=0.006) but not *Rab38* (p=0.70) significantly decreased the expression of *Lamp1*, while the knock down of *Rab29* had the opposite effect on *Lamp1* expression (p=0.004) (**Fig.1 Ce**).

HRP-conjugated mouse monoclonal antibodies raised against Rab32 and Rab38 gave similar increases in the expression of Rab32 and Rab38 with knock down of *Rab29* **(Fig.S1A).** Thus, as a complimentary approach, B16 cells grown on coverslips for knock down of *Rab32*, *Rab38*, *Rab29* or a non-targeting control (NTC) were processed for immunocytochemistry (ICC) and confocal microscopy **(Fig.S1B & C)**. The sub-cellular distribution of Rab29, Rab32 and Rab38 is quite distinct although close homologs. Consistent with published literature (*8*), most endogenous Rab29 was perinuclear, consistent with Golgi localization, and signal intensity diminished with *Rab29* siRNA. Confocal microscopy of endogenous Rab38 displayed a more diffuse albeit discrete, punctate endosomal staining, and that also responded to siRNA-mediated knock down. In contrast, endogenous Rab32 displayed an exclusive, perinuclear punctate staining that was subsequently confirmed to be a Lamp1^+^ compartment, potentially lysosomes **(Fig.S1B, C & D)**.

To further validate the relationship between Rab32, Lamp1 and pRab10, B16 cells knocked down for *Rab32*, *Rab38*, *Rab29* or a NTC were processed for ICC and confocal microscopy. Consistent with our biochemistry experiments, siRNA targeting of *Rab32* significantly reduced the fluorescence intensity of the total Rab32 (p<0.0001) and Lamp1^+^ lysosomes (p=0.001), along with their co-localization (p=0.001), and independent of any difference in pT73-Rab10 (p=0.07), when compared to NTC siRNA. Knock down of *Rab38* had a modest inverse effect on the fluorescence intensity of total Rab32 (p=0.04) and Lamp1 (p=0.08) without significant differences in pT73-Rab10 (p=0.43), while siRNA targeting of *Rab29* induced a significant increase in Rab32 (p<0.0001), Lamp1 (p<0.0001) and pT73-Rab10 (p=0.0002) along with their co-localization (p=0.004; **Fig.1D** & **Ea-d)**. Additionally, confocal microscopy of B16 cells knocked down for Rab32, Rab38 or NTC supported our biochemical studies with the loss of either *Rab32* or *Rab38* having no impact on the fluorescence intensity of endogenous Rab29 **(Fig.S2A & B;** p=0.13 & 0.10, respectively). Together, these data validate the murine Rab29, Rab32 and Rab38 antibodies used, and uncover a novel relationship between the levels of Rab29, Rab32 and Rab38 and their ability to promote Rab10 phosphorylation.

### Rab32 is upstream of Rab38 and Rab29 and is necessary to support Lamp1 and Rab10 phosphorylation

To better understand the functional relationship between Rab29, Rab32, Rab38 and their influence on pT73-Rab10 and Lamp1, B16 cells were plated in siRNA-containing medium against *Rab32*, *Rab38*, *Rab29* or a non-targeting control (NTC) or equimolar concentrations of *Rab38* and *Rab29, Rab32* and *Rab29* or *Rab32* and *Rab38* for 72 hours before western blot (**Fig.S3**). Consistent with the individual targeting siRNA experiments, the co-knock down of Rab38 and Rab29 did not significantly alter the level of Rab32 (**Fig.S3Ba;** siRNA Rab38/29 p=0.06). Similarly, co-knockdown of Rab38 and Rab32 did not significantly affect level of Rab29 (**Fig.S3Bb;** siRNA Rab38/32 p=0.99). Rab38 levels were significantly increased following the knock down of Rab29 (p=0.0001), however this increase was blocked upon co-knock down of Rab29 and Rab32 (**Fig.S3Bc;** p=0.99). There were no significant differences in the level of total Lrrk2 with any of these siRNA treatments (**Fig.S3Bd;** p>0.05). In contrast, while the knock down of Rab29 lead to a significant increase in pT73-Rab10/Rab10 levels (p< 0.0001), this increase was diminished upon co-knock down of Rab38 and Rab29 (p=0.99), Rab32 and Rab29 (p=0.99), and Rab38 and Rab32 (p=0.48) (**Fig.S3Be**). Lamp1 levels were significantly increased following the co-knock down of Rab38 and Rab29 (p=0.002); the co-knock down of Rab32 and Rab29 normalized (p=0.38) the statistically increased levels of Lamp1 observed following the knock down of Rab29 alone (p<0.0001) while the co-knock down of Rab32 with Rab38 significantly reduced the levels of Lamp1 (p=0.04) (**Fig.S3Bf)**. Together, these results suggest that Rab32 is upstream of Rab38 and necessary to promote Rab10 phosphorylation and Lamp1 biogenesis.

### Peripheral administration of LPS increases Rab32 expression and Lrrk2 kinase activity

Recent studies suggest *LRRK2* and *RAB32* expression are immunoresponsive, and that RAB32 promotes LRRK2 kinase activity (*16, 10*). To establish a robust model of inflammation, three-month-old male C57BL/6J WT mice were injected with a single dose of LPS at 5mg/kg, or two single doses of 1mg/kg 24 hours apart, with saline as a control (**Fig.S4**). Brains were harvested at 6 or 24 hours after the last injection. When assessed by qPCR, all mice that received LPS had a increase in the expression of *Irg1*, a known immune-responsive gene/protein that is upregulated during inflammation to produce an anti-microbial metabolite (**Fig.S4Ba-c;** Treatment 1mg/kg, p=0.001 & 0.68, 6 & 24hr, respectively; Treatment 5mg/kg, p=0.001 & 0.04, 6 & 24hr, respectively). However, only mice that received two doses of 1mg/kg LPS had a robust increase in the expression of *Iba1* (Treatment 1mg/kg, p = 0.03 & 0.03, 6 & 24hr) and *Gfap* (Treatment 1mg/kg, p=0.07 & 0.60, 6 & 24hr), which are well described markers of gliosis. To compliment these results, tissues from LPS-injected mice were homogenized and analyzed by western blot. Relative to mice that received saline control, and using 2x 1mg/kg LPS, the levels of Iba1 and Stat1 were significantly increased by 24 hours (Iba1, p=0.002; Stat1, p=0.02) but not at the 6-hour time point (Iba1, p=0.37; Stat1, p=0.68). Mice that received a single dose of LPS at 5mg/kg responded similarly, as indicated by their weight loss (**Fig.S4A**). However, levels of Iba1 and Stat1 within the CNS were unaltered following a single dose of LPS (**Fig.S4B-D**). These findings suggest immune priming using two successive LPS applications is necessary to induce robust CNS inflammation. As a control we analyzed neuronal markers, namely tau and α-synuclein, which remained unaltered by exposure to LPS. Overall, these results suggest that delivery of 2x 1mg/kg of LPS via intraperitoneal injection is sufficient to drive acute inflammation within the CNS without compromising neuronal integrity (**Fig.S4A-D;** p>0.05).

Having established a working inflammation paradigm, we asked whether peripheral inflammation contributes to the expression of Rab32 and Lrrk2 kinase activity *in vivo* within the CNS. Initially, we focused on the midbrain of LPS and saline injected mice as gliosis and nigral neuronal loss are pathologic hallmarks of PD. To ensure that our peripheral inflammatory paradigm was sufficient to induce CNS inflammation, and to validate previously published observations (*16*), we performed qPCR on the midbrain of WT mice that received LPS (2x 1mg/kg at baseline 0, 6 and 24 hr). As expected, the relative gene expression for *Irg1* rapidly increased within 6 hours following LPS exposure (p = 0.04) and throughout the brain but reverted to near saline levels within 24 hours of LPS exposure (**Fig.2A, Fig.S5; Table S2**). Similar to *Irg1,* expression of the transcription factor *Stat1* rapidly increased within 6 hours of LPS exposure (p=0.02) but reverted to near saline levels by 24 hours (p=0.99) (**Fig.2A).** *Iba1* and *Gfap* also increased following peripheral injection with LPS (Iba1, p=0.02 & 0.02, 6 & 24hr; Gfap, p=0.03 & 0.35, 6 & 24hr). Within other sections of the brain, upregulation of *Gfap* was significant in the prefrontal cortex (p=0.0007 & 0.02, 6 & 24hr) and striatum (p<0.0001 & 0.005, 6 & 24hr) but not in the lower brainstem (p=0.99 & 0.50, 6 & 24hr), whereas *Iba1* was especially elevated in the striatum (p=0.034 & 0.023, 6 & 24hr) and lower brainstem (p = 0.08 & 0.02, 6 & 24hr) (**Fig.2A and Fig.S5; Table S2**). Gfap and Iba1 protein levels in the midbrain were significantly elevated by 24 hours (Iba1, 24hr, p=0.003; Gfap, 24hr, p=0.001) (**Fig.2C**). We next assessed the expression profiles of *Rab20* (a published positive control (*16*))*, Rab32, Rab38, Rab29* and *Lrrk2* in the midbrain of saline and LPS injected mice. The relative gene expression of *Rab20* modestly increased at 6 hours following LPS exposure, with the greatest change in the midbrain (p=0.004) and striatum (p=0.04) (**Fig.2, Fig.S5; Table S2)**. *Rab20* gene expression in the midbrain reverted to the level of saline control within 24 hours (p=0.23) (**Fig.S5; Table S2**) although the protein remained elevated (**Fig.2C**; p=0.02, 24hr). The relative gene expression of *Rab32* was significantly increased throughout the brain within 6 hours of LPS exposure (**Fig.S5; Table S2)** and remained elevated in the striatum (p=0.01) and hippocampus (p=0.03) at 24 hours (**Fig.S5; Table S2**), and in midbrain the corresponding levels of protein were significantly elevated (**Fig.2C**; p=0.003). In contrast, *Rab29 and Rab38* gene levels were not significantly increased by LPS-treatment in any brain region (Rab29, p=0.06 & 0.27, 6 & 24hr; Rab38, p=0.70 & 0.22, 6 & 24hr) **(Fig.2, Fig.S5; Table S2**); midbrain Rab29 protein levels were comparable to the saline control at both 6 (p=0.69) and 24 hours (p=0.47), while Rab38 protein levels appeared marginally, albeit non-significantly, elevated (**Fig.2C**; p=0.99 & 0.11, 6 & 24hr). *Lrrk2* gene expression following LPS exposure remained unchanged at 6 and 24 hours (p=0.19 & 0.78, 6 & 24hr), and protein levels were also unchanged (p=0.42 & 0.21, 6 & 24hr) (**Fig.2C and Fig.S5**). Together, these results demonstrate peripheral administration of LPS is sufficient to drive the selective and sustained expression of *Rab32* in midbrain of WT mice. The log2 fold change (FC) of relative gene expression (2^(-ΔΔCt)^) post-LPS for each gene has been mapped to a sagittal ideogram of the mouse brain (**Fig.3**, ideogram courtesy of the GENESTAT Project, and **Fig.S6**). Comparative statistical analysis of gene expression with/without peripheral LPS treatment (6hr and 24hr) within all 6 brain regions is also provided by two-way ANOVA (**Table S2)**.

**Fig. 2.**
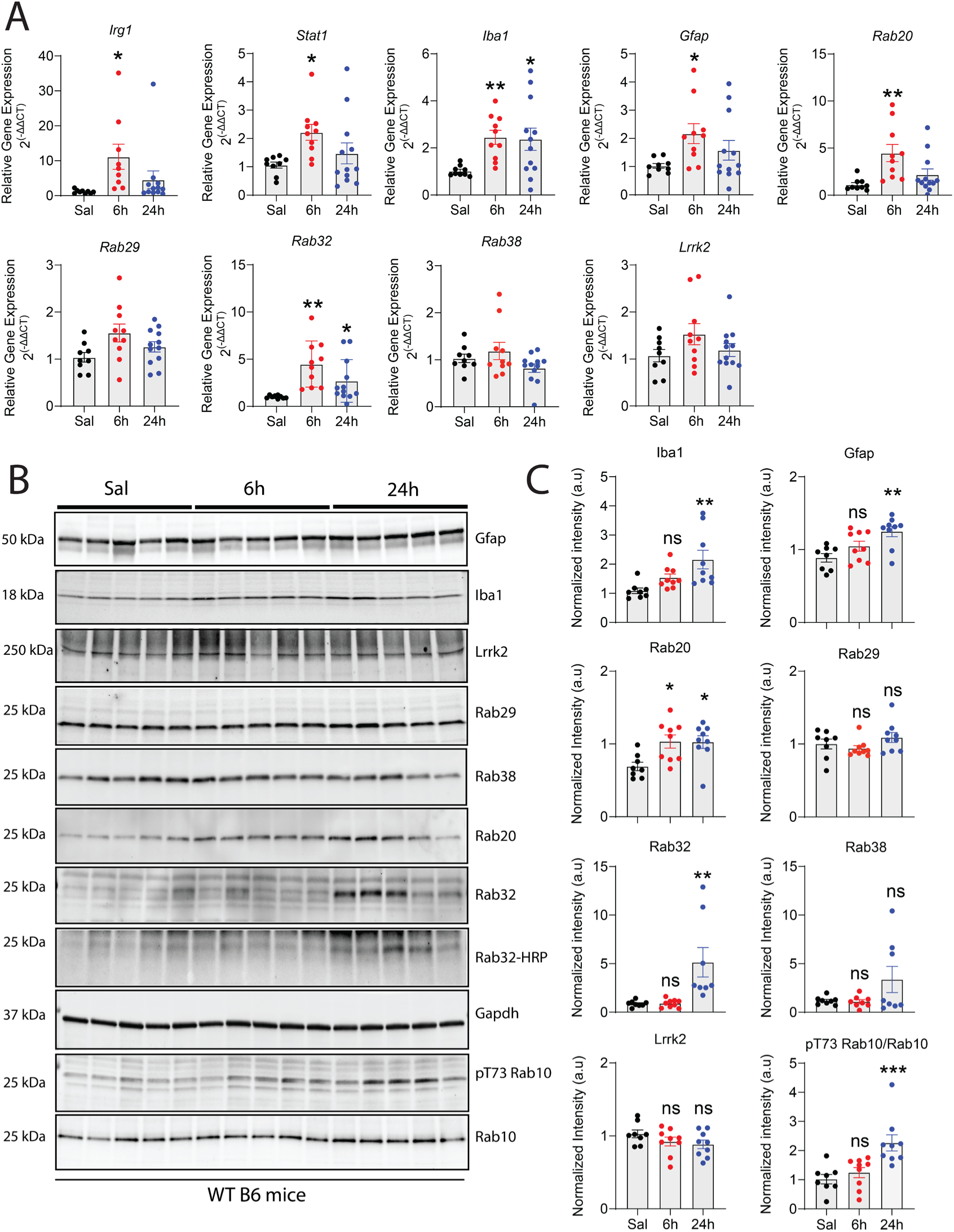
Peripheral Administration of LPS Upregulates Expression of Rab32 and Increases Rab10 Phosphorylation in the Midbrain of WT Mice. **A)** Relative gene expression (2^(-ΔΔCT)^) of in the midbrain of WT mice assessed by qPCR at 6 and 24hours post-injection with 1 mg/kg of LPS or saline control. Each dot on the graph represents an individual animal that underwent the treatment paradigm. **B)** Western blot analysis of RIPA-soluble midbrain lysates at 6 and 24hours post-injection with 1 mg/kg of LPS or saline control. Approximately 10-20ug of protein was resolved for each sample, except for the detection of pT73-Rab10, which required 40ug. **C)** Quantification of western blots shown in **(B)** normalized to total Rab10 or Gapdh loading control. Each dot on the graph represents an individual animal that underwent the treatment paradigm. All data are presented as mean ± S.E.M. Statistical analysis was determined through a one-way ANOVA with a Tukey multiple comparisons post-hoc test, where significant findings are reported relative to NTC. *\*p* <0.05, *\*\*p*<0.01, *\*\*\*p*<0.001.

**Fig. 3.**
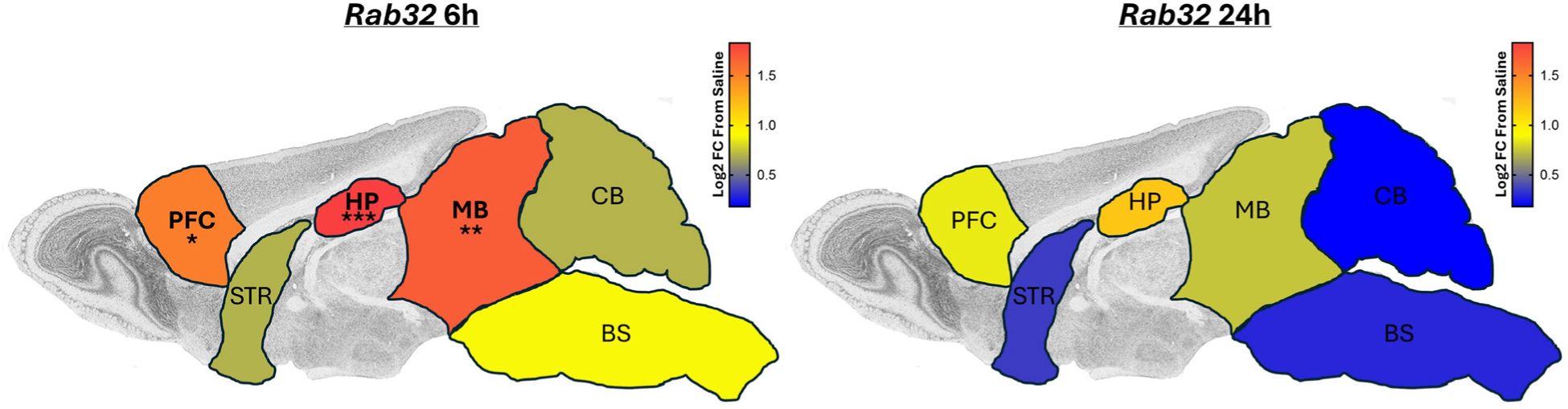
Sagittal ideogram of regional gene expression (2(-ΔΔCT)) of *Rab32* in the brain of wild type mice with LPS. Relative gene expression (2^(-ΔΔCT)^) of *Rab32* from the prefrontal cortex (PFC), striatum (STR), hippocampus (HP), midbrain (MB), cerebellum (CB) and brain stem (BS) of WT mice were assessed by qPCR at 6 and 24 hours post-injection with 1 mg/kg of peripheral LPS or saline control. The log2 fold change (FC) of 2^(-ΔΔCT)^ was transposed as a heatmap onto a sagittal mouse brain ideogram from the GENESTAT Project (https://gensat.org/) (see **Fig. S6** for original brain slice images used). Statistical analysis was determined through a two-way ANOVA with a Tukey multiple comparisons post-hoc test, and significant findings are reported by stars drawn on the brain region. *\*p* <0.05, *\*\*p*<0.01, *\*\*\*p*<0.001. Statistical analysis of gene expression with/without peripheral LPS treatment within all 6 brain regions is by two-way ANOVA (**Table S2)**. The relative gene expression of each individual gene examined within each brain region by one-way ANOVA (**Fig. S5)**.

Next, we assessed the levels of phosphorylated Rab10 (Thr73) normalized to total Rab10 protein, which is an accepted marker of Lrrk2 kinase activity (*17*). Phosphorylated Thr73-Rab10 was significantly increased (p=0.0009) 24 hours after LPS injections, relative to saline control (**Fig.2C**). Phosphorylated Thr73-Rab10 levels were highly correlated with Rab32 levels (F1,24=20.85, r^2^=0.47, p<0.0001, n=26), similar to the correlation for midbrain protein expression of Iba1 (F1,24=19.14, r^2^=0.44, p<0.0002, n=26). Hence, peripheral administration of LPS upregulates inflammatory markers within the CNS of WT mice, including GTPase Rab32 that promotes Lrrk2 kinase activity.

### Peripheral administration of LPS increases Rab32 expression in microglia

Basal Rab32 expression is highest in immune cells, including microglia, according to publicly available datasets (Human Protein Atlas; proteinatlas.org). Thus, we quantified the levels of Rab32 and Lamp1 in Iba1^+^ microglia and Th^+^ DA neurons in sagittal brain sections from WT mice injected with LPS using immunohistochemical methods. Peripheral administration of LPS induced microgliosis, as indicated by the retraction of microglia processes and enlarged cell bodies (**Fig.4A**), and elevated total Rab32 immunoreactivity in the midbrain (**Fig.4B, D**), complementary to biochemical results. In addition, total Lamp1 immunoreactivity was comparable between saline and LPS-injected mice (**Fig.4B**). Interestingly, microglial Lamp1 (p<0.0001) and Rab32 (p=0.0007) were all elevated with LPS, as well as the levels of Rab32 on Lamp1^+^ puncta (p=0.0049) (**Fig.4B**). Conversely, the levels of Lamp1 (p=0.64) and Rab32 (p=0.07) were unaltered with LPS in Th^+^ DA neurons (**Fig.4C, D**).

**Fig. 4.**
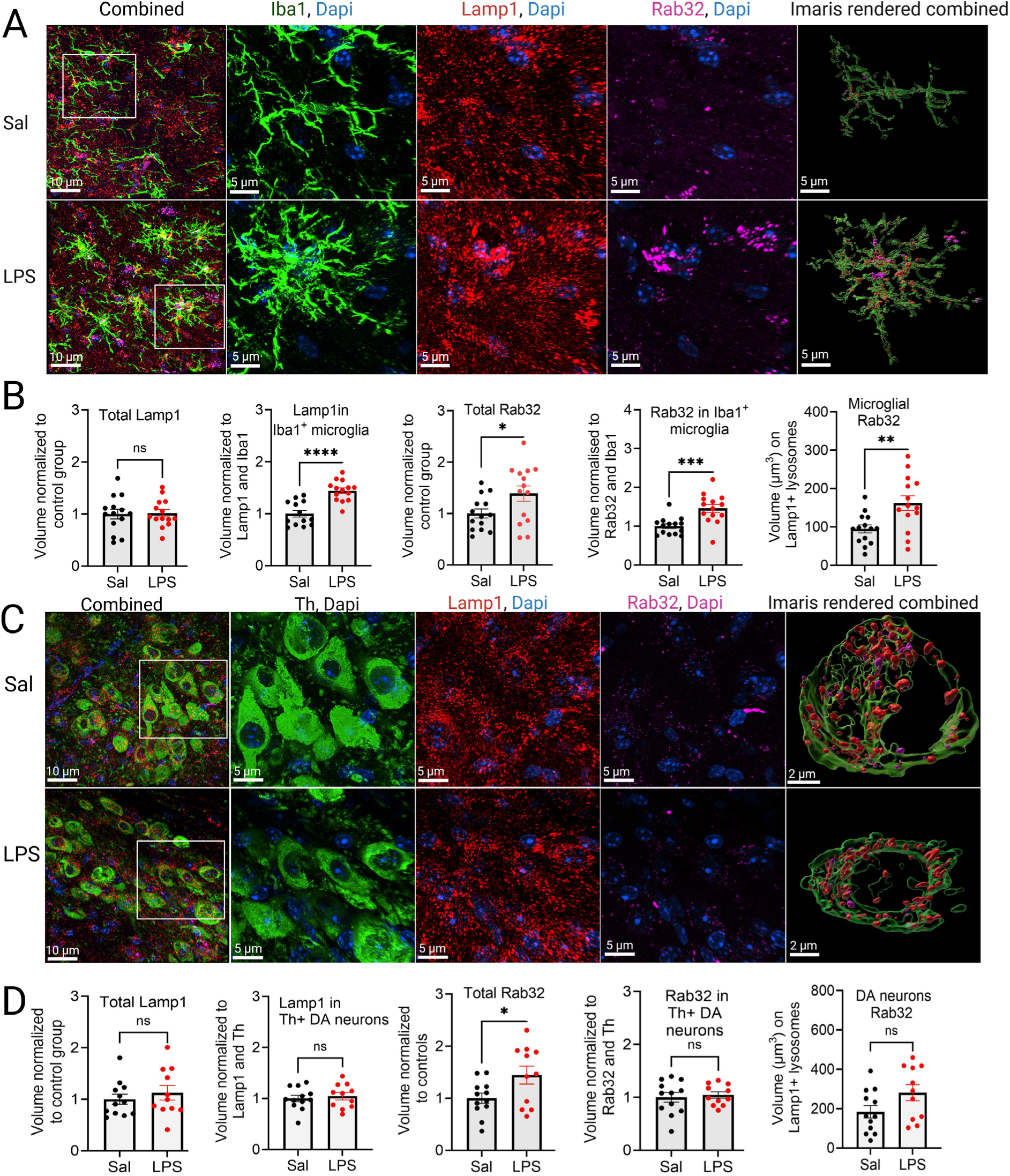
Peripheral administration of LPS in WT mice upregulates expression of Rab32 in microglia but not DA neurons in midbrain. **A)** Representative confocal images of Iba1, Lamp1, Rab32, and their Imaris render, in the midbrain of mice injected with either LPS or saline. Sections are counterstained with DAPI. **B)** Statistical analysis compared saline and LPS treatments. Volumetric analysis of total Lamp1+ puncta; unpaired *t*-test, t26=0.14, p=0.89. Volumetric analysis of Lamp1^+^ puncta within Iba1^+^ microglia; unpaired *t*-test, t25=5.32, p<0.0001. Volumetric analysis of total Rab32^+^ puncta; unpaired *t*-test, t26=2.22, p=0.035. Volumetric analysis of Rab32^+^ puncta within Iba1^+^ microglia; unpaired *t*-test, t26=3.85, p=0.0007. Raw volumetric analysis of microglial Rab32 localized to their Lamp1^+^ lysosomes; unpaired *t*-test, t26=3.08, p=0.0049. **C)** Representative confocal images of Th, Lamp1, Rab32, and their Imaris render, in the midbrain of mice injected with either LPS or saline. Sections are counterstained with DAPI. **D)** Statistical analysis compared saline and LPS treatments. Volumetric analysis of total Lamp1^+^ puncta; unpaired *t*-test, t21=0.75, p=0.46. Volumetric analysis of Lamp1^+^ puncta within DA^+^ neurons; unpaired *t*-test, t21=0.48, p=0.64. Volumetric analysis of total Rab32^+^ puncta; unpaired *t*-test, t21=2.29, p=0.034. Volumetric analysis of Rab32^+^ puncta within in DA^+^ neurons; unpaired *t*-test, t21=0.39, p=0.70). Raw volumetric analysis of DA neuron restricted Rab32 localized to their lamp1^+^ lysosomes; Unpaired *t*-test, t21=1.90, p=0.07). *\*p* <0.05, *\*\*p*<0.01, *\*\*\*p*<0.001, \**\*\*\*p*<0.0001. Error bars are ±SEM. n refers to the number of mice and in parentheses the number of sections [3 M old mice, Sal *n*=7 (14); LPS *n*=7 (14)].

### Nuclear translocation of Tfe3 controls Rab32 expression

*In silico* analysis of murine *Rab32* and human *RAB32* promoters reveal tandem palindromic motifs that approximate CLEAR consensus sequences. These include a 5’ E-box motif (CANNTG), ∼110bp upstream of the transcription start site (TSS), optimal for TFE3 recognition (**Fig.5A; Fig.S8**). *RAB29* and *RAB38* show less conserved CLEAR motifs, none in tandem or near the TSS, consistent with the promoters of their murine homologs. To evaluate the functional relevance of these promoter sequences, we repeated siRNA-mediated knock down of *Rab32*, *Rab38* or *Rab29* in B16 cells and first assessed whether *Tfe3* or *Tfeb* gene expression was altered. While the silencing of *Rab32* or *Rab38* had no detectable effect on *Tfe3* transcript levels when compared to NTC, *Rab29* knock down resulted in modest but significant increase in *Tfe3* expression (p=0.02) (**Fig.5Ba**). In contrast, the knock down of *Rab32* or *Rab29* had no effect on *Tfeb* expression while *Rab38* knock down caused a small reduction in *Tfeb* expression (p=0.01) (**Fig.5Bb)**. Next, we determined whether the increase in *Tfe3* expression observed following *Rab29* knock down was restricted to transcriptional upregulation or accompanied by enhanced nuclear localization of Tfe3 protein. Seventy-two hours after siRNA transfection, B16 cells knocked down for either *Rab32*, *Rab38*, *Rab29* or a non-targeting control (NTC) were collected and processed to obtain crude nuclear and cytoplasmic fractions, or for ICC. Equal amounts of nuclear and cytoplasmic protein were resolved, and fraction purity was confirmed through enrichment of Gapdh or Histone H3, respectively. The competitive mTORC inhibitor Torin was used as a positive control and induced significant nuclear translocation of Tfe3 (p<0.0001) and Tfeb (p<0.0001) and a corresponding >70% dephosphorylation of pSer2448-mTOR/mTOR levels, confirming effective pathway inhibition (p=0.0006). Treatment with Torin had no significant impact on Rab32 (p=0.48) or Lamp1 (p=0.99) protein levels (**Fig.5C & Da)**. Knock down of *Rab32, Rab38* or Rab29 had no significant impact on pSer2448-mTOR/mTOR levels (p=0.37, p=0.25 and p=0.97, respectively). However, quantification of the nuclear-to-cytoplasmic ratio revealed that knock down of *Rab29* (p=0.003), but not *Rab32* (p=0.99) or *Rab38* (p=0.84), caused a significant increase in Tfe3 nuclear translocation **(Fig.5C & Db)**. In contrast, depletion of *Rab32, Rab38,* or *Rab29* had minimal impact on the nuclear-to-cytoplasmic ratio of Tfeb **(Fig. 5C & Dc**; p=0.99, p=0.99 and p=0.08, respectively). To complement, B16 cells grown on coverslips knocked down for *Rab32, Rab38* or *Rab29* were fixed and co-stained for endogenous Rab32 and Tfe3. Consistent with biochemical results, short-term inhibition of the mTOR1 complex with Torin induced a robust translocation of endogenous Tfe3 from the cytoplasm to the nucleus (p < 0.0001) **(Fig.5E & F)**. In contrast to the knock down of *Rab32* or *Rab38*, the loss of *Rab29* induced a statistically significant increase in the nuclear-to-cytoplasmic ratio of *Tfe3* (p = 0.003) **(Fig.5E & F)**. Taken together, these results suggest that the loss of *Rab29* expression promotes a distinct Tfe3-Rab32 pathway that occurs independent of gross mTOR inhibition.

**Fig. 5.**
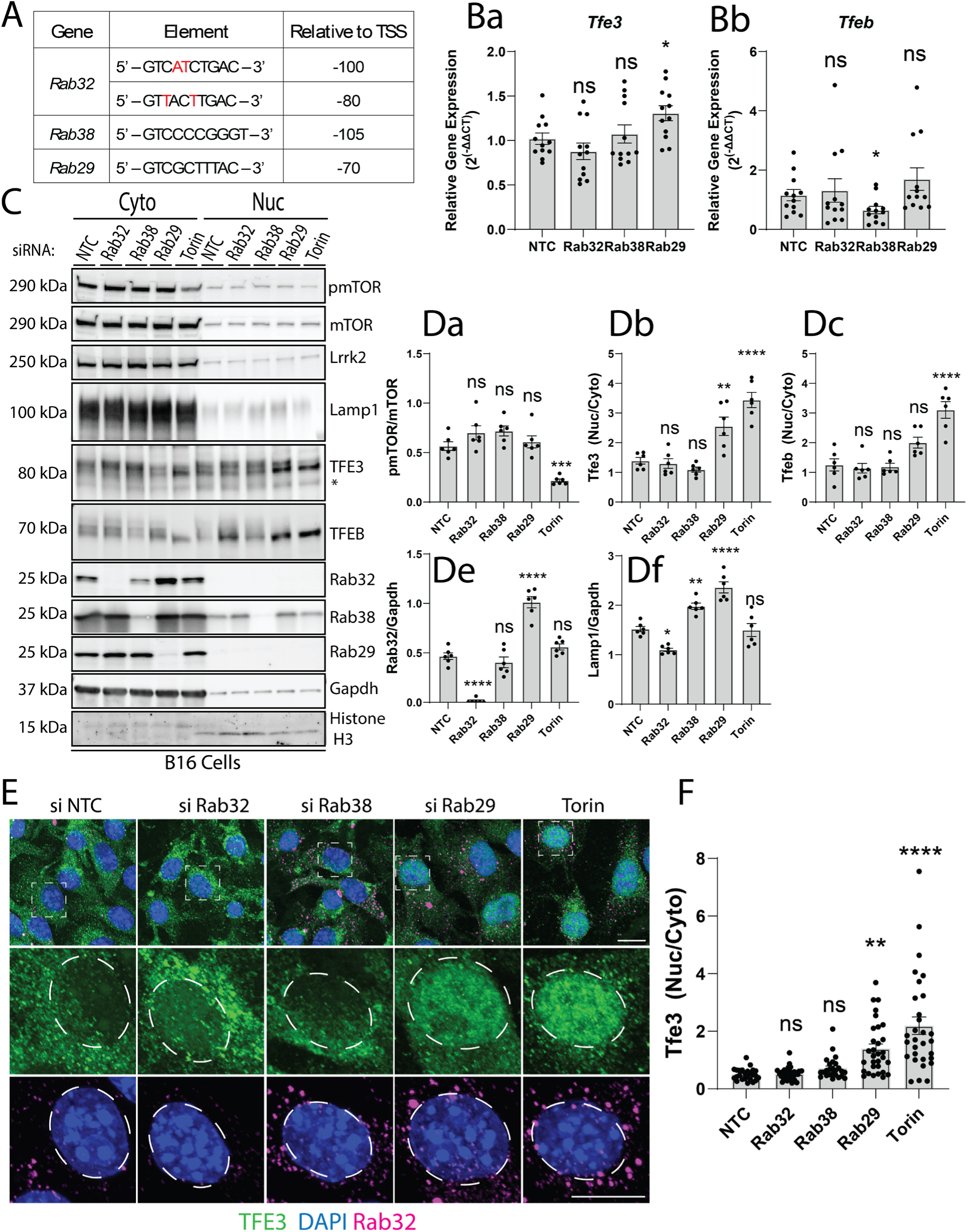
Loss of Rab29 expression promotes nuclear translocation of Tfe3 and controls Rab32 expression. **A)** Position of sequences that approximate a palindromic 10-bp GTCACGTGAC CLEAR motif, albeit with 2 or 3 bp mismatches. In Rab32, but not Rab38 or Rab29, these are found in tandem and include an E-box (CANNTG) (see Fig.S8). **B)** Relative gene expression (2(-ΔΔCT)) of Tfe3 and Tfeb expression in B16 cells following siRNA-mediated knock down of Rab32, Rab38 or Rab29. Data is pooled from six independent experiments where each dot on the graph represents an individual 3 combined wells that underwent the treatment paradigm. **C)** Representative western blot data of cytosolic and nuclear fractions from B16 cells knock down for Rab32-sub family of GTPases. Data is representative of six independent experiments. * denotes non-specific Tfe3 band. **D)** Quantification of protein lysates analyzed in (C) **E)** Representative confocal microscopy of Tfe3 (green) and Rab32 (magenta) in B16 cells following siRNA-mediated knock down. Nuclei are counterstained with DAPI. Oval ROI denotes representative nuclear perimeter used for analysis. Scale bar = 10µm. **F)** Quantification of nuclear-to-cytoplasmic fluorescence intensity of Tfe3 presented in (E). Data presented as mean ±S.E.M. Statistical analysis was determined through a one-way ANOVA with a Tukey multiple comparisons post-hoc test, where significant findings are reported relative to NTC. *\*p* <0.05, *\*\*p*<0.01, *\*\*\*p*<0.001, \**\*\*\*p*<0.0001.

### Nuclear translocation of Tfe3 and increased Rab32 expression in Iba1+ microglia following exposure to LPS

Previous literature suggests Tfeb, alongside Tfe3, may work in unison to regulate lysosomal biogenesis, cytokine production and immune cell function in response to inflammation. Owing to our data in B16 cells implicating Tfe3 and not Tfeb in regulating Rab32 expression, we assessed the expression and localization of *Tfeb* and *Tfe3* in the midbrain of WT mice using our established model of inflammation in three-month-old male C57BL/6J WT mice (**Fig.2 & Fig.S4**). When assessed by qPCR, all mice that received LPS had a significant increase in the levels of *Tfe3* (p=0.007), but not *Tfeb* (p=0.33) at 6 hours post LPS injection that returned to near-baseline levels within 24 hours (**Fig.6Aa & Ab;** Tfe3, p=0.08; Tfeb, p=0.99). Immunohistochemistry and confocal microscopy revealed an increased proportion of Iba1^+^ microglia exhibiting nuclear Tfe3 (**Fig.6B & Ca;** p=0.009), a greater relative volume of Tfe3 within Iba1^+^ microglia (**Fig.6B & Cb;** p=0.0012), and a concomitant shift in the nuclear-to-cytoplasmic localization of Tfe3 **(Fig.6B & Cc;** p<0.0001). To validate these findings and extend our observations to a human-relevant cellular model, we generated iPSC-derived microglia from healthy donors. Following treatment with LPS (100ng/mL; 24 hours) or saline control, iPSC-derived microglia were fixed and stained with antibodies against Rab32 and Iba1. Iba1 immunoreactivity was observed in >95.0% of cells across both treatment conditions but exposure to LPS induced a more polarized microglia phenotype. Compared with saline control, LPS treatment significantly increased total Rab32 immunoreactivity (p=0.0023) (**Fig.6Da & Db**) and promoted nuclear translocation of endogenous Tfe3 (p<0.0001) (**Fig.6Ea & Eb**) in iPSC-derived microglia. These results suggest that LPS induces Rab32 expression in murine and human iPSC-derived microglia by promoting the translocation of Tfe3 to the nucleus.

**Fig. 6.**
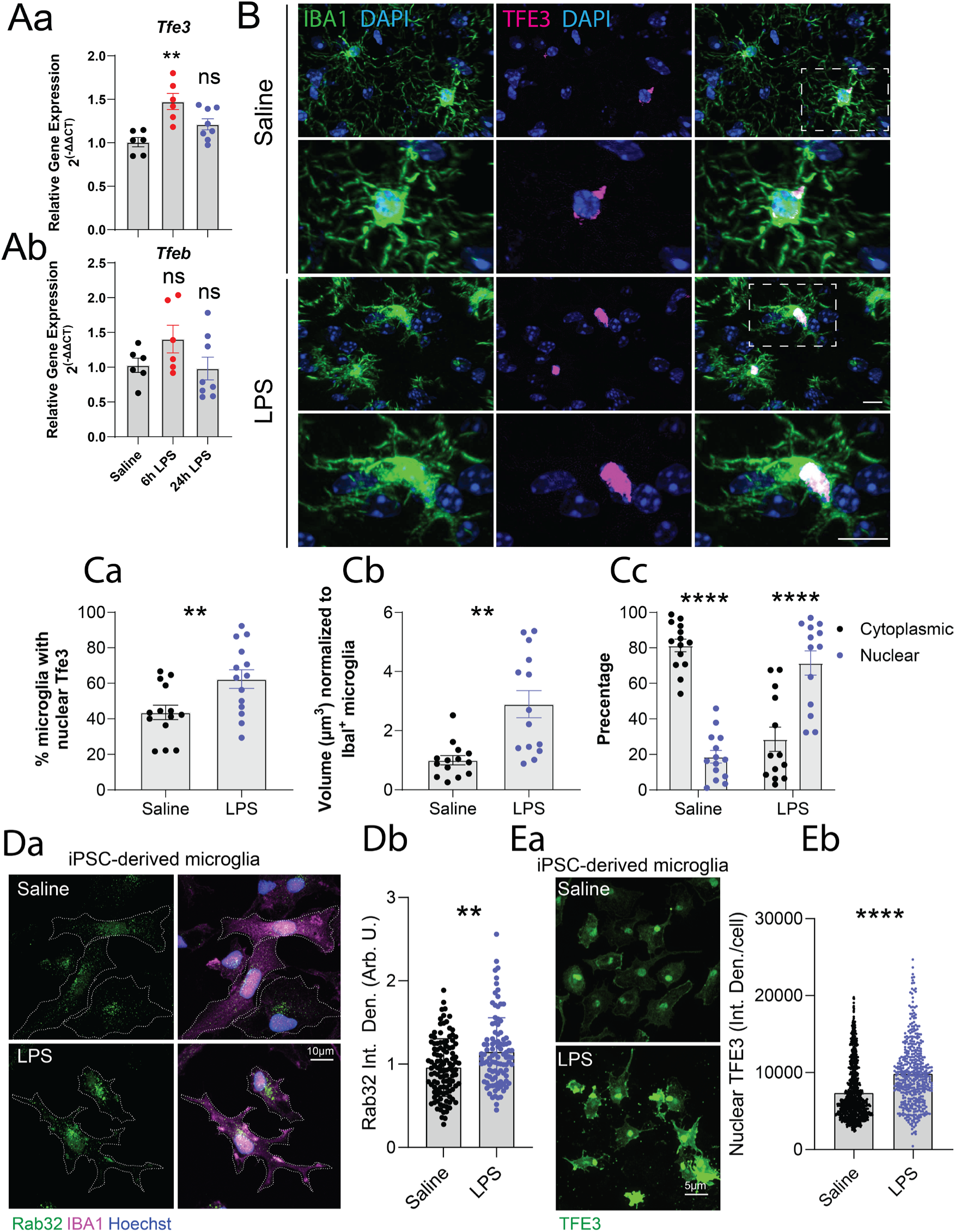
Microglial TFE3 is immune responsive and translocate to the nucleus upon exposure to LPS. **A)** Relative gene expression (2(-ΔΔCT)) of *Tfe3* and *Tfeb* in the midbrain of WT mice assessed by qPCR at 6 and 24 hours post-injection with 1 mg/kg of LPS or saline control. Each dot on the graph represents an individual animal that underwent the treatment paradigm. Data presented as mean ± S.E.M. **B)** Representative confocal images of Iba1 and Tfe3 in the midbrain of mice injected with either LPS or saline. Sections are counterstained with DAPI. **C)** Quantified percentage of microglia with nuclear Tfe3 staining presented in **B.** Statistical analysis was assessed by an unpaired *t*-test, t26=2.82, p=0.009. **Cb)** Volumetric analysis of Tfe3 within Iba1^+^ microglia. Statistical analysis was assessed by an unpaired *t*-test with Welch’s correction, t15.9=3.93, p=0.0012. **Cc)** Relative percentage of cytoplasmic vs. nuclear Tfe3 within Iba1^+^ microglia following treatment with LPS. **D)** Representative confocal microscopy images of endogenous Rab32 and IBA1 in IPSC-derived microglia treated with 100ng/mL LPS or saline control. Cells are counterstained with DAPI. **Db)** Quantification of fluorescence intensity for microscopy images presented in **Da**. Data is representative of > 100 cells quantified across 3-independent experiments. Error bars mean ±SEM. **E)** Representative confocal microscopy images of endogenous Tfe3 IPSC-derived microglia treated with 100ng/mL LPS or saline control. **Eb)** Quantification of fluorescence intensity for microscopy images presented in **Ea**. Data is representative of >100 cells quantified across 3-independent experiments. Error bars mean ±SEM. Statistical analysis was determined through an unpaired *t*-test, \**\*\*\*p*<0.0001. All data is presented as mean ±S.E.M. For Fig. 6A, statistical analysis was assessed by a Welch and Brown-Forsythe one-way ANOVA with Dunnett’s multiple comparison testing. For **Fig.6Ca,** an unpaired *t*-test was used; Fig. 6Cb, Welsch’s *t*-test was used while a two-way ANOVA with Šídák’s multiple comparisons test was used for Fig. 6Cc; *\*p* <0.05, *\*\*p*<0.01, *\*\*p*<0.001, *\*\*\*p*<0.0001. Error bars are ± SEM. n refers to the number of mice and in parentheses the number of sections [3 M old mice, Sal n = 7 (14); LPS n = 7 (14)]. Fig. 6Db Statistical analysis was determined through a Mann-Whitney *t*-test while Fig. 6 **Eb** used an unpaired *t*-test.

### Knock down of Tfe3 mitigates increased Rab32 and pRab10 phosphorylation

To validate our observation that Tfe3 localization regulates Rab32 expression and link this pathway with Lrrk2 activation, we performed concurrent knock down of *Rab32*, *Rab38*, *Rab29* or a non-targeting control (NTC) together with knock down of *Tfeb* or *Tfe3* for 72 hours, then processed samples for western blot. Under these experimental conditions, the expression of Tfe3 and Tfeb appear inter-dependent, consistent with their established function as a heterodimer.

Furthermore, the steady-state levels of Tfe3, but not Tfeb, in buffer containing 1% triton, a non-ionic detergent that solubilizes lipid membranes without disrupting the nucleus, were inversely correlated with the levels of Rab32 (Tfe3, p<0.0001; Tfeb, p=0.82) and Rab38 (Tfe3, p<0.0001; Tfeb, p>0.99) while the knock down of Rab29 had a modest yet statistically significant reduction in Tfe3 levels (p=0.0004) without impacting Tfeb (p=0.99)(**Fig.7Aa & Ab)**, likely reflecting its nuclear translocation (**Fig.5C & Db)**. In contrast, the knock down of *Rab32*, *Rab38*, *Rab29* or a non-targeting control (NTC) together with knock down of *Tfeb* or *Tfe3* had no effect on the steady-state levels of Lrrk2, Rab38 or Rab29 (**Fig.7Ac & Ae**). While the knock down of *Tfe3* or *Tfeb* on either a NTC or *Rab38* down background had minimal effect on steady-state levels of Rab32, pT73-Rab10/Rab10 or Lamp1, *Tfe3 -* but not *Tfeb* – knock down on a Rab29 knock down background restored the statistically significant increase in Rab32, pT73-Rab10/Rab10 and Lamp1 to near-control levels (**Fig.7A & Ba - Bc**). Lastly, we employed confocal microscopy of endogenous Rab32, pT73-Rab10 and Lamp1 in B16 cells concurrently knocked down for Rab29 or a non-targeting control (NTC) together with knock down of Tfe3 or Tfeb. Consistent with our biochemical results, the statistically significant increase in Rab32, pRab10 and Lamp1 puncta observed upon the loss of Rab29 were restored upon silencing of *Tfe3* and not *Tfeb*. Together, these results identify Rab32 as a predominant driver of Lrrk2 activation and demonstrate that expression is regulated through Tfe3 nuclear localization.

**Fig. 7.**
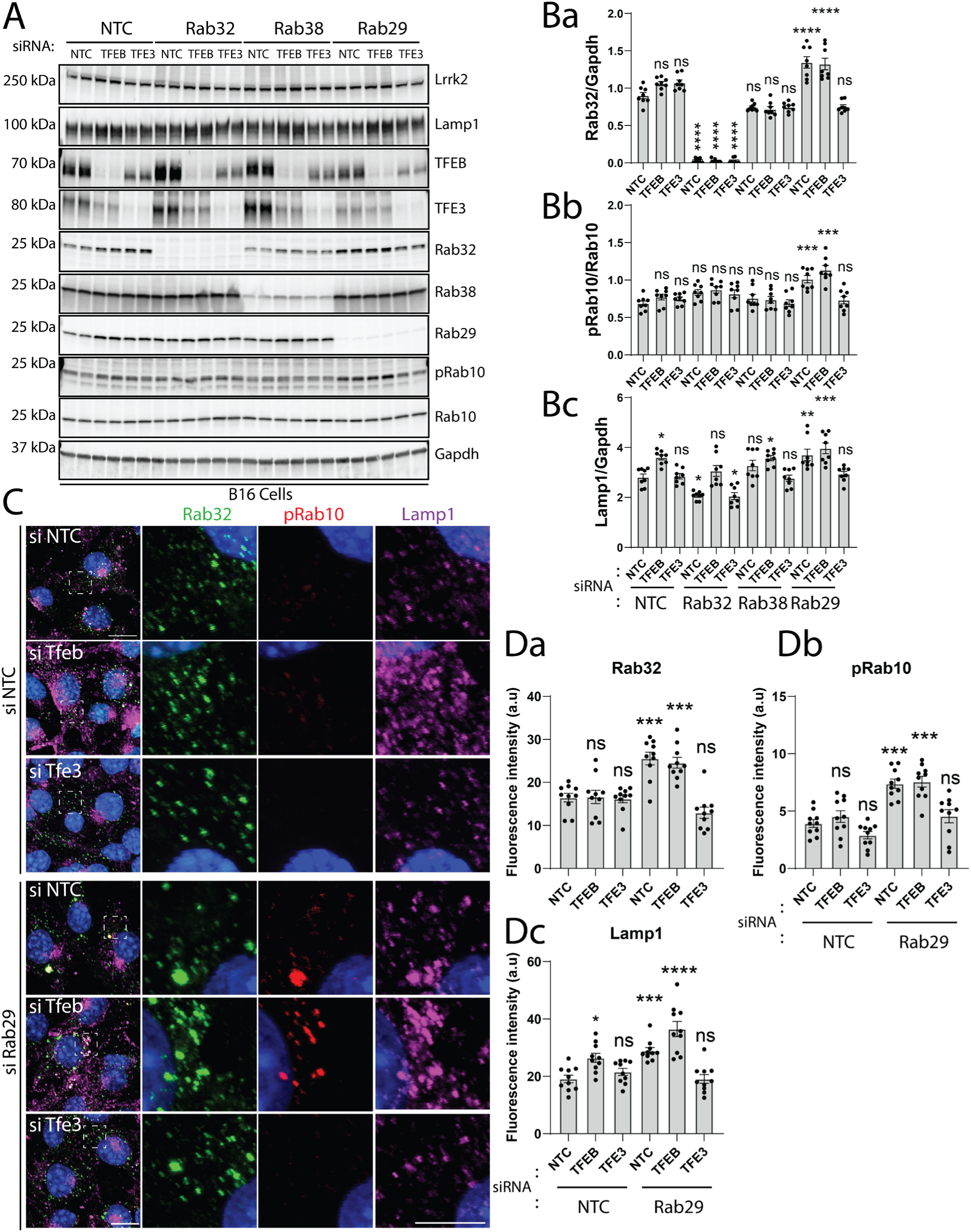
Knock down of Tfe3 mitigates increased expression of Rab32 and pRab10 phosphorylation. **A)** Representative western blot data of Rab32-sub family of GTPases, Lrrk2, Lamp1 and pT73-Rab10/Rab10 in siRNA *Tfe3* or *Tfeb* knock down B16 cells. Data is representative of four independent experiments. **B)** Quantification of protein lysates analyzed in **(A)**. **C)** Representative confocal microscopy of Rab32, pT73-Rab10 and Lamp1 in B16 cells following the indicated siRNA-mediated co-knock down. Nuclei are counterstained with DAPI. Images are from two independent experiments with 4-7 images captured per experiment. **D)** Quantification of fluorescence intensity analyzed in **(C)** All data presented as mean ±S.E.M. Statistical analysis was determined through a two-way ANOVA with a Tukey multiple comparisons post-hoc test, where significant findings are reported relative to NTC. *\*p* <0.05, *\*\*p*<0.01, *\*\*\*p*<0.001, \**\*\*\*p*<0.0001.

## DISCUSSION

Recent studies suggest RAB29 is the main GTPase responsible for LRRK2 recruitment to membrane for kinase activation (*8, 6, 18*). In part, this originated from a modest genetic association between common variants in the *RAB29* locus and PD (*8*), which remains robust (rs11557080 A>G odds ratio (OR)=1.12, 95% CI 1.09-1.16, p=1.85E-11)(*5*). Nevertheless, Lrrk2 kinase activity is not attenuated in *Rab29* germline knock out mice (*9*). Recent genetic data now links its homolog, RAB32 Ser71Arg as a Mendelian gene for PD and, by association this mutation has a much greater effect on the likelihood of PD (OR=13.17, 95% CI 2·15–87·23, p=0.0055) leading to a 2-fold increase in LRRK2 kinase activity (*10*). Rare variants in *RAB29* and *RAB38* that activate LRRK2 kinase have yet to be linked or associated with PD. Here we demonstrate RAB32 specifically localizes to a Lamp1^+^ compartment, in contrast to RAB29 and RAB38, consistent with the major role of LRRK2 in endolysosomal autophagy. Notably, our *in vitro* results suggest that transiently lowering *Rab29* leads to a modest compensatory increase in Rab32 or Rab38 protein expression, but not vice versa, with analogous findings by gene expression. Silencing *Rab29* also leads to a significant increase of pThr73-Rab10/Rab10 arguing against its necessity for physiological Lrrk2 membrane recruitment and kinase activation. The amount of Lamp1 protein also significantly increases when *Rab29* or *Rab38* are silenced but marginally diminishes when *Rab32* is silenced, suggesting a dependency on the latter and, intriguingly, the transcription levels of Lamp1 appear to mediate this effect.

Rab32 is the only Group II GTPase with immune-responsive expression. *In vivo* peripheral LPS results in almost immediate transcriptional upregulation of inflammatory markers in the brain, including *Iba1*, *Gfap*, *Rab20* and *Rab32* within 6 hours of last administration, of which *Iba1* and *Rab32* levels remain elevated at 24 hours, in close agreement with prior results (*16*). Corresponding midbrain protein levels appear to be highly correlated albeit temporally delayed, with Iba1, Stat1, Rab20 and Rab32, all significantly elevated 24 hours post LPS exposure.

Although Lrrk2 protein levels did not change with this stimulus or over this time course, Lrrk2 kinase activity (measured as the ratio of pThr73-Rab10/Rab10) was also highly correlated with the elevation of Rab32 (r^2^=0.46, p<0.0001). Additional regional analysis of gene expression demonstrated Rab32 was upregulated in the lower brainstem, midbrain, hippocampus and prefrontal cortex within 6 hours, and maintained its elevated expression at 24 hours. How closely this effect may be associated with LPS dosing, and how long it lasts after administration requires further study.

*In vitro* cell culture and immunoprecipitation studies demonstrate Lrrk2 kinase activity requires membrane recruitment. All three homologs, RAB29, 32 and 38, specifically bind LRRK2 through its ankyrin domain and can recruit LRRK2 monomer to membrane. Nevertheless, each of these RAB GTPases localizes to different intracellular compartments namely the Golgi, endoplasmic reticulum or Lamp1^+^ vesicles, potentially lysosomes, respectively. Whether one specific membrane interaction is more relevant for the pathogenesis of PD remains unclear. *In vivo*, in the adult mouse brain and under non-inflammatory conditions, Iba^+^ microglia typically have a small cell body with highly ramified branches. Here peripheral LPS treatment appears to induce a hyper-ramified state associated with Rab32 and Lamp1 upregulation. Complimentary results, most notably an increased expression of Rab32 following acute treatment with LPS, were observed in iPSC-derived microglia (**Fig. 6**). In midbrain microglia there was also a highly significant co-localization of these markers on Lamp1^+^ vesicles that was concomitantly associated with increased Lrrk2 kinase activity. In contrast, neither Rab32 nor Lamp1 within dopaminergic neurons were significantly increased with LPS treatment. Activated microglia are known to mediate non-cell-autonomous inhibition of neuronal autophagy via paracrine signaling (*19*) but how Rab32 levels influences cytokine secretion in the peripheral immune system and microglia has yet to be established (*20, 21*).

Only RAB32 has tandem CLEAR sequences ∼110bp proximal to its TSS, consistent with optimal TFE3 transcription factor recognition. Both TFEB and TFE3 transcription factors are associated with lysosomes in a phosphorylated state, in part, mediated by RAB32, mTOR activity and nutrient sensing to downregulate autophagy (*22*). However, infection in myeloid-cells increases TFEB and TFE3 binding to CLEAR motifs located in the promoters of many lysosomal and proinflammatory genes to increase their expression (*23, 24*). LRRK2 kinase activity has been shown to modulate the expression and nuclear localization of TFE3 to inhibit lysosomal biogenesis (*25*), whereas mitochondrial translocation of TFEB also regulates complex I and inflammation (*26*). Now our results suggest RAB32 expression, likely in response to TFE3-induction and nuclear localization in microglia, provides a reciprocal mechanism to increase LRRK2 kinase activity in a circuit we term the “RAB32 rheostat” (**Fig. 5, 6, 7**). In myeloid-derived cells this mechanism helps regulate autophagosomal capacity for pathogen clearance by macrophages, allergic response by mast cells, and controls antigen presentation by dendritic cells and immune signaling (*27*). However, to understand the roles of RAB32 and LRRK2 hyperactivity on immune surveillance, synaptic pruning and phagocytosis by microglia, and on dopaminergic neuronal survival in PD, demands further studies in model systems based on this insight.

From these data, we also postulate reduced/incomplete penetrance in individuals and families with *LRRK2* mutations (*28, 29*) is a function of *RAB32* expression driven by intermittent peripheral immune activation throughout their lifespan. Such a hypothesis is consistent with PD as a multifactorial disease, manifest with age and inflammatory exposures, and may provide an elegant molecular explanation for the body-first hypothesis of PD (*30*). It would also be consistent with the inverse association between the use of non-steroidal anti-inflammatory medications in *LRRK2* PD (*31*). Population differences in pathogen exposure may explain the disparity in disease penetrance in LRRK2 Gly2019Ser heterozygotes in Tunisia versus Norway, given that the cumulative incidence of idiopathic PD with age is almost identical (*29*). The inter-individual variance in *RAB32* expression in human subjects has yet to be assessed but may be an important consideration for the success of clinical trials of Lrrk2 kinase inhibitors. The GTEx Portal (gtexportal.org) shows many single nucleotide polymorphisms in the *RAB32* locus are correlated with its expression in different tissues, including peripheral nerve and brain, and further studies of related genetic and biomarker variation in multi-incident *LRRK2* pedigrees may inform disease penetrance.

*RAB32* biology provides an unequivocal physiologic and molecular conduit that unites research observations in PD including its clinical presentation, epidemiologic risk factors and prior Mendelian gene discoveries. This thesis can be viewed from an evolutionary genetic perspective, driven by the relationship between humans and pathogens (*3*). RAB32 expression, LRRK2 kinase activity, TFEB/TFE3 phosphorylation and subsequent translocation to the nucleus or mitochondria appears to be intimately controlled from the lysosomal compartment to modulate peripheral and central immune responses, metabolism and dopaminergic cell survival. This nexus warrants future investment to develop therapeutic interventions in PD that offer more than symptomatic benefit.

## MATERIALS AND METHODS

### Animals and housing

All animal procedures were performed in accordance with a protocol approved by the Institutional Animal Care and Use Committee (IACUC) at the University of Florida. Colonies were maintained on a reverse cycle (light on from 7pm to 7am) and single-sex group-housed in enrichment cages after weaning at post-natal day 21. All mice were supplied with rodent diet and water *ad libitum*. All experiments, data processing and analysis were conducted in a blinded manner. All mice utilized had a C57BL/6J congenic background from Jackson Labs.

### Cell lines, siRNA treatments, RNA and protein extraction

Mouse melanoma B16-F10 (CRL-6475; ATCC©) cells were maintained at 37 °C/5% CO2 in Dulbecco’s Modified Eagles Medium (DMEM) from ATCC© (Cat #30-2002) and Thermo Fisher Scientific© (Cat #11965118), respectively, supplemented with 10% (v/v) FBS (Gibco©; 16140071) and 2mM Glutamax (Gibco©: 35050061). Cells were passaged every 2-3 days, and passage number was maintained below 20 to limit cellular senescence.

For siRNA experiments, B16 cells were seeded in DMEM growth medium containing 25 nM pooled siRNA complexes in Nunc 6-well culture dishes (Thermo Fisher Scientific©; Cat # 140685) at 1 ×10^6^ per well. siRNA complexes against *Rab32 (*Catalog ID: L-063539-01-0020), *Rab38* (Catalog ID: L-040873-01-0020*), Rab29* (Catalog ID: L-053101-01-0020), *Tfeb (*ID:L-050607-02-0020), *Tfe3* (ID:L-054750-00-0020) and non-targeting control (Catalog ID:D-001810-10-20) were purchased from Dharmacon (Lafayette, CO, USA).

After seventy-two hours of transfection, cells were processed for RNA extraction using a RNeasy® Mini Kit (Qiagen©) or protein extraction using ice-cold 1x Cell Signaling Lysis Buffer (Cell Signaling Technology© #9803S) supplemented with Halt phosphatase inhibitor cocktail (Thermo Fisher Scientific©), and protease inhibitor cocktail (Roche©) and left on ice for 30 min to lyse. Lysates were clarified at 20,000 g for 10 min at 4 °C and pelleted debris were removed.

Samples were supplemented with NuPage LDS sample buffer 4x (NP0008), boiled for 5 min at 95 °C and analyzed by standard western blot protocols.

### Peripheral injection of lipopolysaccharide (LPS) in C57BL/6J mice

LPS was purchased from Cell Signaling Technology (Cat # 14011) and reconstituted in sterile PBS to a working stock of 1mg/ml. To obtain volumes suitable for intraperitoneal injection (i.p.) LPS stock solution was diluted in sterile PBS to a concentration of 0.5mg/ml or 0.1mg/ml, filtered with Millex filter unit (0.22 um) and injected in sterile glass vials to be used during injection to limit contamination. Mice were injected with a single dose of 5mg/kg of LPS, or 2 subsequent doses of 1mg/kg LPS (24 hr apart), then euthanized for tissue collection via inhalation of isoflurane at 6 and 24 hr after the last dose. Control mice were given saline.

### Brain extraction and homogenization for RNA and protein extraction

Mice were euthanized by isoflurane inhalation and transcardially perfused with ice-cold phosphate-buffered saline (PBS). Harvested brains were generally midline dissected with one hemisphere drop-fixed in 4% paraformaldehyde (PFA) in PBS for 24 hr for subsequent immunofluorescence (IF) while cortex (CTX) and midbrain (MB) were micro-dissected from the other hemisphere, and snap-frozen in liquid nitrogen for quantitative polymerase chain reaction (qPCR) and/or western blotting. From a separate cohort, brains were cut at the midbrain into left and right hemispheres, and micro-dissected into prefrontal cortex (PFC), striatum (STR), hippocampus (HP), midbrain (MB), lower brainstem (BS; both pons and medulla) and cerebellum (CB) to examine regional gene expression of immunoregulation and Rab GTPase genes. Snap-frozen brain sections designated for RNA extraction were first mechanically homogenized in RLT buffer (provided in the RNeasy® Mini Kit [Qiagen©]) with a glass mortar and was further homogenized using QIAshredder (Qiagen©) tubes. After homogenization, brain tissues underwent RNA extraction/isolation following the protocols provided by the RNeasy® Mini Kit, where homogenates were treated with 70% ethanol and RNA was captured and washed within the mini spin columns provided. Pure RNA was eluted into separate 1.5 microcentrifuge tubes from the mini columns with 50 μL of RNase-free water (provided by the RNeasy® Mini Kit) and each sample was then quantified (ng/uL) using a NanoDrop™ One/One^C^ spectrophotometer (Thermo Fisher Scientific©). All RNA samples were kept in aliquots frozen at -80 °C until ready for use. Brain tissues designated for protein extractions were homogenized in RIPA Lysis Buffer (Pierce® # 89900) supplemented with Halt™ Protease and Phosphatase Inhibitor Cocktail (100x; Thermo-Fisher™: 78440) in a 1 ml glass Dounce homogenizer (Wheaton) and allowed to rotate at 4°C for 45 min on HulaMixer before centrifugation (14,000 × *g* at 4 °C for 15 min). Supernatant was aliquoted appropriately to help mitigate freeze-thawing cycles. Homogenate aliquots and the resulting pellets were frozen separately and stored at -80 °C.

### Single strand complementary DNA (cDNA) synthesis and quantitative PCR

RNA samples from cohorts of mice and B16 cell lines were converted into cDNA with a High-Capacity cDNA Reverse Transcription Kit (Applied Biosystems™). Briefly, in a 384-well plate, a mixture containing 10 μL of total cDNA master mix (10X RT Buffer, 10X RT Random Primers, 25x dNTP Mix [100 mM], MultiScribe™ Reverse Transcriptase [50 U/μL]) and 10 μL of 400 ng of RNA (diluted with RNase-free water) were dispensed across three wells per mouse/cell line per gene examined (3 technical replicates). The 384-well plate was then sealed with optical tape and placed in a VeritiPro™ 384-well Thermal Cycler (Applied Biosystems™) for reverse transcription. After cDNA conversion, 4 μL from each well were pipetted into a new 384-well plate, where 10 μL of TaqMan™ Universal Master Mix II with uracil-N-glycosylase (Applied Biosystems™), 5 μL of RNase/DNase-free water, and 1 μL of TaqMan™ Gene Expression Assay (Thermo Fisher Scientific©) was added in three wells per mouse/B16 cell line, per gene examined. TaqMan™ Gene Expression Assays used were: glyceraldehyde-3-phosophate dehydrogenase (*Gapdh*): Mm99999915_g1; beta-actin (*Actb*): Mm02619580_g1; Ionized calcium binding adaptor molecule 1 (*Aif1/Iba1*): Mm00479862_g1; glial fibrillary acidic protein (*Gfap)*: Mm01253033_m1; immune-response gene 1 (*Irg1*): Mm01224532_m1; Lysosome-associated membrane protein 1 (*Lamp1*): Mm01217070_m1; Signal transducer and activator of transcription 1 (*Stat1*): Mm01257286_m1; *Lrrk2*: Mm00481934_m1; *Rab20*: Mm01216807_m1; *Rab29*: Mm00523103_m1; *Rab32*: Mm00458960_m1; *Rab38*: Mm00503697_m1. Once cDNA and TaqMan™ master mixes were pipetted, the 384-well plates were placed into a QuantStudio™ 5 Real-Time PCR System (Applied Biosystems™) using software to conduct qPCR and find comparative Ct values (ΔΔCt) from each well. For each mouse/B16 cell line examined on the qPCR plates, the combined mean of Ct values from the housekeeping genes *Gapdh* and *Actb* were calculated and ΔCt was found for each targeted gene from each sample (ΔCt = Ct value of targeted gene – Ct value of combined mean *Gapdh* and *Actb*). To compare groups from saline controls (mice cohorts) and no template controls [NTC] (B16 cell line), the ΔΔCt method was used, where ΔΔCt = ΔCt of targeted gene from experimental group – ΔCt of targeted gene from control group. Once ΔΔCt was found for each comparison, relative gene expression for the gene of interested in each sample was calculated using 2^(-ΔΔCt)^.

### Nuclear and cytoplasmic protein isolations, and western blotting

B16 cells seeded in Nunc 6-well dishes underwent nuclear and cytoplasmic extraction using the NUC-PER kit (Thermo Fisher Scientific; Cat # 78833). Three hours prior to collection, cell culture medium was replenished on all wells, at which point Torin (1µM; Cell Signaling; 14379) was added as a positive control for nuclear translocation. All buffers were supplemented with protease and phosphatase inhibitors (Thermo Fisher Scientific Cat # 78441) prior to use.

Protein samples were subjected to Pierce™ BCA Protein Assay Kit (Thermo Scientific™), protein concentrations were calculated and samples reduced with NuPage™ LDS Sample Buffer 4x (#NP0008) for 5 min at 95 °C. Samples were then analyzed by standard western blotting using primary antibodies detailed in **Table S1**. Secondary antibodies were used at 1:10,000 dilution: IRDyes® 800CW Goat anti-Rabbit IgG (Li-Cor Biotech©; #926–32211) and 680RD Goat anti-Mouse IgG (Li-Cor Biotech©; #926–68070). Raw blots were quantified using ImageJ’s western blotting algorithm to determine the stained area under the curve and normalized to the levels of Gapdh.

### Immunofluorescence, immunohistochemistry and confocal microscopy

B16 cells were seeded on GelTrex (Gibco) coated 12 mm glass coverslips in siRNA-containing DMEM. After 72 hours, cells were washed twice in RT PBS, then fixed in 4% PFA/PBS for 15 mins at RT. Fixed cells were washed in 1 x PBS (3 x 5 mins), permeabilized with 0.1% Triton-X100/PBS (v/v) at RT for 12 mins, blocked for 45 minutes with 5% NGS/PBS (v/v) and then incubated overnight at 4°C with primary antibodies listed in **Supplemental Table 1**, diluted in 2% NGS/PBS/0.02% Tween-20 under constant agitation. After 16 hours, the cells were washed (3 x 5 mins) with 5% NGS/PBS (v/v) before incubation at RT with Alexa Fluor secondary antibodies and DAPI (1:5000) for 90 minutes. Cells were washed (4 x 5 mins) at RT in 1 X PBS before mounting in Fluoromount-G^TM^ (00-4958-02, Invitrogen©).

Brains (24-hour time point only) were fixed in 4% paraformaldehyde (PFA) overnight at 4°C, and cryoprotected with 30% sucrose in PBS before 30 µm sagittal slices were obtained with a cryostat. Sections were blocked in 10% normal goat serum (NGS) in 0.3% PBS-triton-X 100 (PBST) (1 hour, 37°C), followed by overnight (4°C) incubation of primary antibodies, described in **Supplemental Table 1,** diluted in 5% NGS + 0.3% PBST. The sections were washed (4×15 min in PBS), followed by secondary antibody incubation with species-specific Alexa Fluor™ IgG secondary antibodies for 1h at RT (Invitrogen©, 1:500). Tissues were rewashed in PBS (4 × 15 min), then co-stained with DAPI (Thermo Fisher Scientific©, 1:5000) and mounted using Fluoromount-G^TM^ (00-4958-02, Invitrogen©). Images were acquired at 60x field of view on a Nikon® Spinning Disc Confocal Microscope (AX/AX R with NSPARC) using NIS-Elements AR Imaging software from Nikon® (90 z-stack steps at 0.30 μm spacing), with its intrinsic deconvolution function. Th, Iba1, Lamp1 and Rab32 immunoreactivity in the MB were quantified with Imaris^TM^ software (Oxford Instruments). Briefly, Th^+^ DA neurons (smoothing: 0.3 μm) and Iba1^+^ microglia (smoothing; 0.10 μm) were rendered as surfaces, and the AI component of the Imaris^TM^ software was used for unbiased identification of Th^+^ DA neurons and Iba1^+^ microglia - only whole cells in the field of view were included in the analysis as identified with DAPI. The microglia projections that broke off from their somas were stitched, and the volume of Th^+^ DA neurons and Iba1^+^ microglia were quantified accordingly. Similarly, Lamp1 (smoothing: 0.10 μm, threshold value: 500 and number of voxels: 10) and Rab32 puncta (smoothing: 0.10 um, threshold value: 250, and number of voxels: 10) were rendered as surfaces, their volumes were quantified accordingly and normalized to the total volumes of Th and/or Iba1.

### Differentiation, immunofluorescence and imaging of human iPSCs into microglia

Human iPSCs were maintained feeder-free in StemFlex medium (Thermo Fisher) on Matrigel-coated plates and passaged with Accutase. The microglia differentiation protocol was adapted from (*32*). Briefly, 10,000 cells/well were seeded into 96-well ultra-low attachment plates in embryoid body (EB) medium (E8 supplemented with 10 µM ROCK inhibitor, 50 ng/ml BMP-4, 20 ng/ml SCF, 50 ng/ml VEGF-121) and centrifuged (300 g, 3 min). EBs were cultured for 4 days with a half-medium change on day 2. On day 4, EBs were transferred to 6-well plates (20/well) in microglia progenitor medium consisting of X-VIVO 15 media (Lonza; 02-053Q) supplemented with 2 mM GlutaMax, 55 µM β-mercaptoethanol, 100 ng/ml M-CSF, 25 ng/ml IL-3). Medium was refreshed twice weekly. Progenitors appeared between days 7–21 and were harvested from suspension. Progenitors were plated in microglia maturation medium consisting of Advanced RPMI 1640 (Thermo Fisher Scientific) with 2 mM GlutaMax, 100 ng/ml IL-34 (PeproTech), and 10 ng/ml GM-CSF (PeproTech). Cells adhered within 1–2 hours and matured into microglia over 6–10 days. Parent EB cultures continued to produce progenitors for up to 3 months with regular feeding. All cytokines were from PeproTech unless otherwise specified. For LPS treatment experiments, progenitors were plated at 150,000 cells per well in a 24-well plate, and following 6-10 day culture in maturation media, microglia were treated with 100 ng/mL LPS for 24 hours prior to fixation.

### Nomenclature and statistical analysis

By convention all human gene names are italicized and capitalized, mouse gene names only have the first letter capitalized, whereas the protein’s encoded are written as standard text. Relative gene expression in B16 cells, and following siRNA-mediated knock down, was analyzed using a repeated-measures one-way ANOVA with a Tukey’s multiple comparisons *post hoc* test.

Relative gene expression between mice cohorts were analyzed using either Brown-Forsythe and Welch ANOVA tests, with a Dunnett’s T3 multiple comparisons *post hoc* test (for parametric data), or a Kruskal-Wallis test with a Dunn’s multiple comparisons *post hoc* test (for nonparametric data), or a two-way ANOVA with a Tukey multiple comparisons *post-hoc* test.

Figure legends indicate which tests were used. Experiments examining phosphorylation and total protein levels by immunoblotting were analyzed by one- or two-way ANOVA with Tukey’s *post-hoc* test. Immunohistochemical images were analyses via unpaired *t*-test. All statistical analyses were performed using GraphPad Prism for Windows (GraphPad Software, San Diego, CA). Mean values ± S.E.M are presented with each dot on the graph representing an individual experimental unit.

## Acknowledgements

This study was supported in part by research grants from the Michael J. Fox Foundation (MJFF-025964; awarded to MJF), the National Institutes of Health grant AG077269 (A.M.), and the Center for Translational Research in Neurodegenerative Disease (CTRND; awarded to JF and MJF). We thank the animal care staff at the McKnight Brain Institute and the Cancer and Genetics Research Complex, University of Florida, Gainesville, Florida, USA, the CTRND at the University of Florida for its technical support and access to shared equipment and Dr. Dario Alessi and the MRC PPU Reagents and Services at the University of Dundee for contributing Rab32 plasmids to this study. We also thank Dr. John Landers (UMass Chan Medical School, Worcester MA), Dr. Mark Cookson (NIA/NIH, Bethesda, MD) and the iPSC Neurodegenerative Disease Initiative (iNDI) for the iPSC cell lines, protocols and technical support.

## Contributions

J.F.: conceptualization and project design, data curation, formal analysis, writing original draft, review and editing, and funding; I.B.D.: conceptualization and project design, data curation and formal analysis, writing original draft, review and editing; R.C.S.: conceptualization and project design, data curation and formal analysis, writing original draft; review and editing; S.W.: data curation, editing and reviewing; M.J.F.: resources and funding, conceptualization, writing, editing and reviewing.

## Competing interests

The authors declare no competing interests.

---

**Supplementary Table 1:** Antibodies used throughout the study.

| **Antibody** | **Source** | **Species** | **ICC/IHC** | **WB** |
| --- | --- | --- | --- | --- |
| Rab32 | Santa Cruz; sc-390178 | Mouse | 1:100/1:500 | 1:1000 |
| Rab38 | Santa Cruz; sc-390176 | Mouse | 1:200 | 1:1000 |
| Rab29 | Abcam [MJF-R30-124]; ab256526 | Rabbit | 1:200 | 1:2000 |
| Gapdh | Thermo Fisher Scientific; MA5-15738 | Mouse |  | 1:5000 |
| Iba | Synaptic Systems, 234 308 | Guinea pig | 1:500 |  |
| Iba1 | Wako; 016-20001 | Rabbit |  | 1:1000 |
| Stat1 | Cell Signaling; 9172S | Rabbit |  | 1:2000 |
| Lamp1 | Developmental Studies Hybridoma Bank© [DSHB]; 1D4B-c | Rat | 1:300/1:500 |  |
| Lamp1 | Abcam [EPR21026]; ab208943 | Rabbit |  | 1:3000 |
| Th | EnCor Biotechnology©, CPCA-TH | Chicken | 1:2000 |  |
| pThr73 Rab10 | Abcam [MJF-R21];<br>ab230261 | Rabbit |  | 1:1000 |
| Rab10 | Abcam [MJF-R23];<br>ab237703 | Rabbit |  | 1:2000 |
| Lrrk2 | Abcam [MJFF2 c41-2; ab133474, Abcam | Rabbit |  | 1:1000 |
| Tfeb | Cell signaling;<br>#83010 | Rabbit |  | 1:2000 |
| Tfe3 | Cell Signaling;<br>#42733. | Rabbit |  | 1:2000 |
| Tfe3 | Millipore-Sigma<br>(SAB5700877). | Rabbit | 1:200 |  |

## Notes

### Competing Interest Statement

The authors have declared no competing interest.

### Summary of Updates

Revised and updated text and figures.

## REFEREENCES

1. H. Sekiya, L. Franke, Y. Hashimoto, M. Takata, K. Nishida, N. Futamura, K. Hasegawa, H. Kowa, O. A. Ross, P. J. McLean, T. Toda, Z. K. Wszolek, D. W. Dickson, Widespread distribution of α-synuclein oligomers in LRRK2-related Parkinson’s disease. Acta Neuropathol 149, 42 (2025).

2. T. Lüth, B.-H. Laabs, S. Sendel, I. R. König, A. Caliebe, A. J. Noyce, L. A. Screven, S. Bardien, M. Farrer, F. Hentati, R. Amouri, C. Klein, S. B. Sassi, J. Trinh, G. P. G. Program (GP2), The Age at Onset of LRRK2 p.Gly2019Ser Parkinson’s Disease Across Ancestries and Countries of Origin. Annals of Neurology **n/a**, doi:10.1002/ana.78181.

3. D. T. Guenther, J. Follett, R. Amouri, S. B. Sassi, F. Hentati, M. J. Farrer, The Evolution of Genetic Variability at the LRRK2 Locus. Genes (Basel) 15, 878 (2024).

4. E. McGrath, D. Waschbüsch, B. M. Baker, A. R. Khan, LRRK2 binds to the Rab32 subfamily in a GTP-dependent manner via its armadillo domain. Small GTPases 12, 133–146 (2021).

5. T. G. P. G. Program (GP2), H. L. Leonard, Novel Parkinson’s Disease Genetic Risk Factors Within and Across European Populations, 2025.03.14.24319455 (2025).

6. E. G. Vides, A. Adhikari, C. Y. Chiang, P. Lis, E. Purlyte, C. Limouse, J. L. Shumate, E. Spínola-Lasso, H. S. Dhekne, D. R. Alessi, S. R. Pfeffer, A feed-forward pathway drives LRRK2 kinase membrane recruitment and activation. eLife 11 (2022), doi:10.7554/ELIFE.79771.

7. D. R. Alessi, E. Sammler, LRRK2 kinase in Parkinson’s disease. Science 360, 36–37 (2018).

8. D. A. MacLeod, H. Rhinn, T. Kuwahara, A. Zolin, G. Di Paolo, B. D. McCabe, B. D. MacCabe, K. S. Marder, L. S. Honig, L. N. Clark, S. A. Small, A. Abeliovich, RAB7L1 interacts with LRRK2 to modify intraneuronal protein sorting and Parkinson’s disease risk. Neuron 77, 425–39 (2013).

9. A. F. Kalogeropulou, J. B. Freemantle, P. Lis, E. G. Vides, N. K. Polinski, D. R. Alessi, Endogenous Rab29 does not impact basal or stimulated LRRK2 pathway activity. Biochemical Journal 477, 4397–4423 (2020).

10. E. K. Gustavsson, J. Follett, J. Trinh, S. K. Barodia, R. Real, Z. Liu, M. Grant-Peters, J. D. Fox, S. Appel-Cresswell, A. J. Stoessl, A. Rajput, A. H. Rajput, R. Auer, R. Tilney, M. Sturm, T. B. Haack, S. Lesage, C. Tesson, A. Brice, C. Vilariño-Güell, M. Ryten, M. S. Goldberg, A. B. West, M. T. Hu, H. R. Morris, M. Sharma, Z. Gan-Or, B. Samanci, P. Lis, M. T. Periñan, R. Amouri, S. Ben Sassi, F. Hentati, F. Tonelli, D. R. Alessi, M. J. Farrer, RAB32 Ser71Arg in autosomal dominant Parkinson’s disease: linkage, association, and functional analyses. The Lancet Neurology 0 (2024), doi:10.1016/S1474-4422(24)00121-2.

11. H. Lian, D. Park, M. Chen, F. Schueder, M. Lara-Tejero, J. Liu, J. E. Galán, Parkinson’s disease kinase LRRK2 coordinates a cell-intrinsic itaconate-dependent defence pathway against intracellular Salmonella. Nature microbiology 8, 1880–1895 (2023).

12. Z. Wu, H. Que, C. Li, L. Yan, S. Wang, Y. Rong, Rab32 family proteins regulate autophagosomal components recycling. Journal of Cell Biology 223 (2024), doi:10.1083/JCB.202306040/276549.

13. S.-L. Lu, S. Chen, K. Noda, Y. Li, C.-Y. Tsai, H. Omori, Y. Kato, Z. Zhang, B. Chen, K. Tokuda, T. Zheng, M. Wakita, E. Hara, M. Fukuda, Y. Wada, E. Morita, N. Uzawa, S. Murakami, T. Noda, Evidence that mitochondria in macrophages are destroyed by microautophagy. Nat Commun 16, 8123 (2025).

14. M. Sardiello, M. Palmieri, A. di Ronza, D. L. Medina, M. Valenza, V. A. Gennarino, C. Di Malta, F. Donaudy, V. Embrione, R. S. Polishchuk, S. Banfi, G. Parenti, E. Cattaneo, A. Ballabio, A Gene Network Regulating Lysosomal Biogenesis and Function. Science 325, 473–477 (2009).

15. A. Bentley-DeSousa, D. Clegg, S. M. Ferguson, LRRK2, lysosome damage, and Parkinson’s disease. Current Opinion in Cell Biology 93, 102482 (2025).

16. Y. Liang, S. Lin, L. Zou, H. Zhou, J. Zhang, B. Su, Y. Wan, Expression profiling of Rab GTPases reveals the involvement of Rab20 and Rab32 in acute brain inflammation in mice. Neuroscience letters 527, 110–114 (2012).

17. M. Bu, J. Follett, I. Deng, I. Tatarnikov, S. Wall, D. Guenther, M. Maczis, G. Wimsatt, A. Milnerwood, M. S. Moehle, H. Khoshbouei, M. J. Farrer, Inhibition of LRRK2 kinase activity rescues deficits in striatal dopamine physiology in VPS35 p.D620N knock-in mice. npj Parkinson’s Disease 2023 9:1 9, 1–12 (2023).

18. H. Zhu, F. Tonelli, M. Turk, A. Prescott, D. R. Alessi, J. Sun, Rab29-dependent asymmetrical activation of leucine-rich repeat kinase 2. Science 382, 1404–1411 (2023).

19. B. P. Festa, F. H. Siddiqi, M. Jimenez-Sanchez, H. Won, M. Rob, A. Djajadikerta, E. Stamatakou, D. C. Rubinsztein, Microglial-to-neuronal CCR5 signaling regulates autophagy in neurodegeneration. Neuron 111, 2021–2037.e12 (2023).

20. N. Tandon, E. Gutiérrez-Martínez, P. Bourdely, G. Anselmi, J. Denisot, Y. Gerber, J. Helft, M. Bens, L. Saveanu, P. Guermonprez, The Rab32 small GTPase is required for efficient cross-priming of CD8+ T cells against cell-associated antigens by XCR1+ type 1 DCs in vivo, 2025.05.03.652057 (2025).

21. R. Song, C. Ngoka, A. Singla, D. A. Kramer, A. Diwaker, D. J. Boesch, Q. Liu, J. J. Moresco, B. Beutler, H.-C. Reinecker, E. Burstein, B. Chen, E. E. Turer, The Rab32-LRMDA-Retriever Complex is a Key Regulator of Intestinal Immune Homeostasis, 2025.07.16.665158 (2025).

22. X. Li, Y. Chen, S. Gong, H. Chen, H. Liu, X. Li, J. Hao, Emerging roles of TFE3 in metabolic regulation. Cell Death Discov 9, 93 (2023).

23. J. A. Martina, H. I. Diab, O. A. Brady, R. Puertollano, TFEB and TFE3 are novel components of the integrated stress response. EMBO J 35, 479–495 (2016).

24. N. Pastore, A. Vainshtein, T. J. Klisch, A. Armani, T. Huynh, N. J. Herz, E. V. Polishchuk, M. Sandri, A. Ballabio, TFE3 regulates whole-body energy metabolism in cooperation with TFEB. EMBO Molecular Medicine 9, 605–621 (2017).

25. N. Yadavalli, S. M. Ferguson, LRRK2 suppresses lysosome degradative activity in macrophages and microglia through MiT-TFE transcription factor inhibition. Proceedings of the National Academy of Sciences of the United States of America 120 (2023), doi:10.1073/PNAS.2303789120.

26. C. Calabrese, H. Nolte, M. R. Pitman, R. Ganesan, P. Lampe, R. Laboy, R. Ripa, J. Fischer, R. Polara, S. K. Panda, S. Chipurupalli, S. Gutierrez, D. Thomas, S. M. Pitson, A. Antebi, N. Robinson, Mitochondrial translocation of TFEB regulates complex I and inflammation. EMBO Rep 25, 704–724 (2024).

27. J. E. Irazoqui, Key Roles of MiT Transcription Factors in Innate Immunity and Inflammation. Trends Immunol 41, 157–171 (2020).

28. D. G. Healy, M. Falchi, S. S. O’Sullivan, V. Bonifati, A. Durr, S. Bressman, A. Brice, J. Aasly, C. P. Zabetian, S. Goldwurm, J. J. Ferreira, E. Tolosa, D. M. Kay, C. Klein, D. R. Williams, C. Marras, A. E. Lang, Z. K. Wszolek, J. Berciano, A. H. Schapira, T. Lynch, K. P. Bhatia, T. Gasser, A. J. Lees, N. W. Wood, Phenotype, genotype, and worldwide genetic penetrance of LRRK2-associated Parkinson’s disease: a case-control study. The Lancet Neurology 7, 583–590 (2008).

29. F. Hentati, J. Trinh, C. Thompson, E. Nosova, M. J. M. J. Farrer, J. O. J. O. J. O. Aasly, LRRK2 parkinsonism in Tunisia and Norway: A comparative analysis of disease penetrance. Neurology 83, 568–569 (2014).

30. P. Borghammer, The brain-first vs. body-first model of Parkinson’s disease with comparison to alternative models. J Neural Transm (Vienna) 130, 737–753 (2023).

31. M. S. Luciano, C. M. Tanner, C. Meng, C. Marras, S. M. Goldman, A. E. Lang, E. Tolosa, B. Schüle, J. W. Langston, A. Brice, J.-C. Corvol, S. Goldwurm, C. Klein, S. Brockman, D. Berg, K. Brockmann, J. J. Ferreira, M. Tazir, G. D. Mellick, C. M. Sue, K. Hasegawa, E. K. Tan, S. Bressman, R. Saunders-Pullman, NON-STEROIDAL ANTI-INFLAMMATORY USE AND LRRK2 PARKINSON’S DISEASE PENETRANCE. Mov Disord 35, 1755–1764 (2020).

32. P. W. Brownjohn, J. Smith, R. Solanki, E. Lohmann, H. Houlden, J. Hardy, S. Dietmann, F. J. Livesey, Functional Studies of Missense TREM2 Mutations in Human Stem Cell-Derived Microglia. Stem Cell Reports 10, 1294–1307 (2018).

