## Supplemental Figures for "Peripheral inflammation mediates midbrain Lrrk2 kinase activity via Rab32 expression"

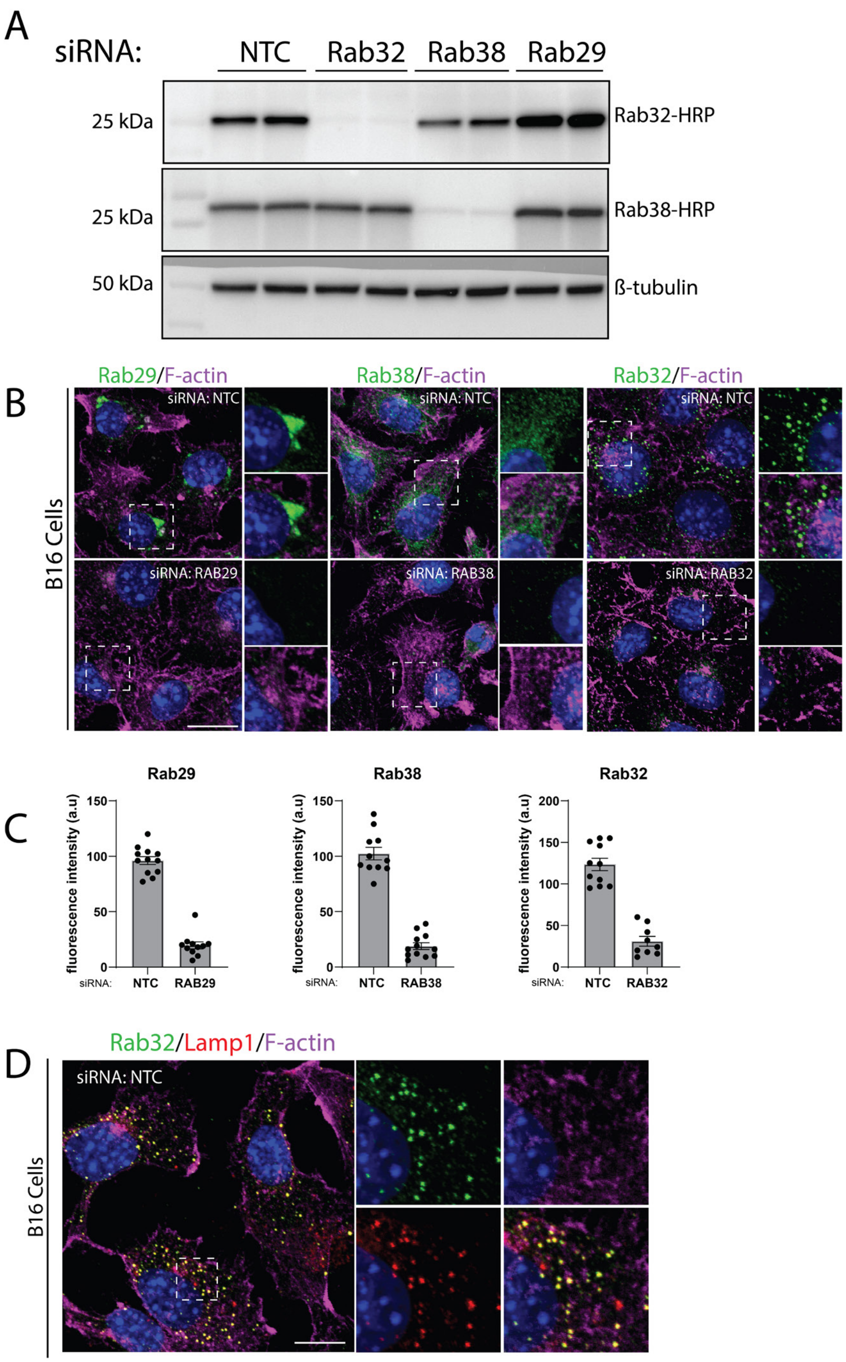


**Fig. S1. Validation of antibodies against Rab32, Rab38 and Rab29 by immunocytochemistry in B16 Cells.**

- **A)** Validation of Rab32 and Rab38 antibody specificity via HRP-conjugated antibodies.
- **B)** Representative confocal microscopy images of endogenous Rab32, Rab38 or Rab29 with and without targeting siRNAs. Cells are counterstained with DAPI. Scale bar = 10µm.
- **C)** Quantification of fluorescence intensity for microscopy images presented in B.
- **D)** Representative confocal image of Rab32 localization to LAMP1$^{+}$ lysosomes in B16 cells. Cells are counterstained with DAPI.

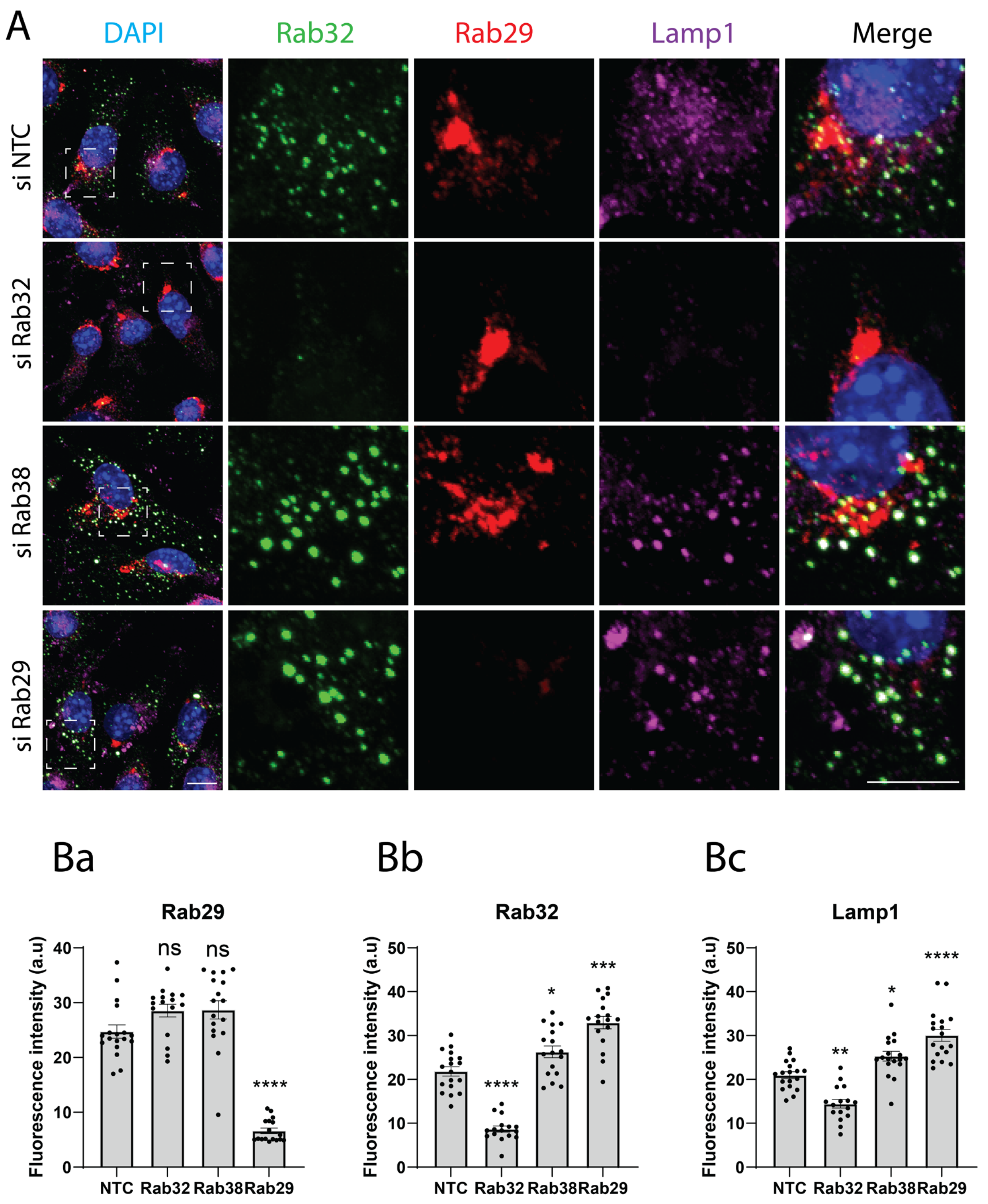


**Fig. S2. Knock down of Rab32 or Rab32 in B16 cells does not impact Rab29 localization.**

- **A)** Representative confocal microscopy images of endogenous Rab32, Rab29 and Lamp1 with and without targeting siRNAs. Cells are counterstained with DAPI. Scale bar = 10µm.
- **B)** Quantification of fluorescence intensity analyzed in (**A**) Data presented as mean ±S.E.M. Statistical analysis was determined through a one-way ANOVA with a Tukey multiple comparisons post-hoc test, where significant findings are reported relative to NTC. *$p$ <0.05, **$p$<0.01, ***$p$<0.001, ****$p$<0.0001.

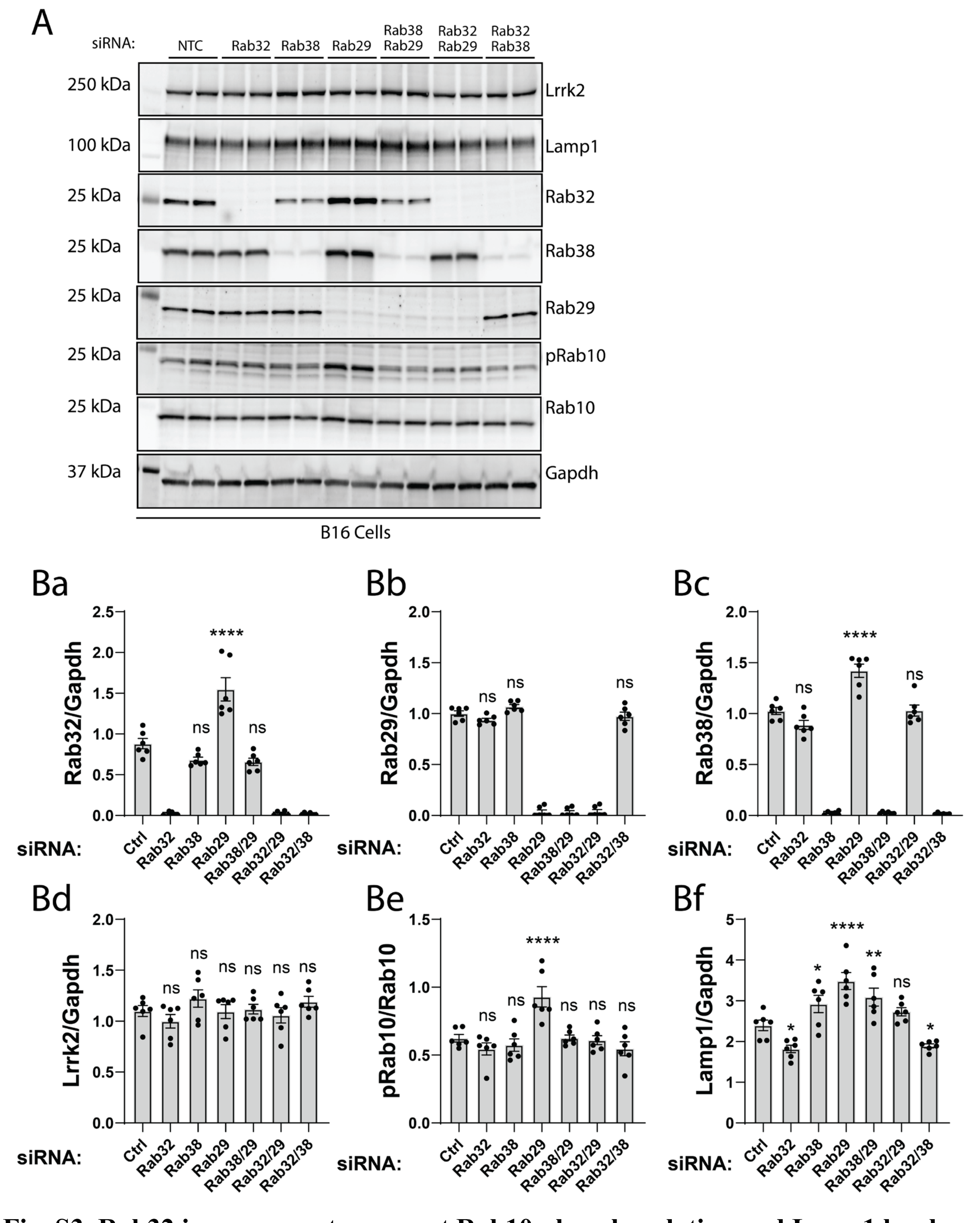


**Fig. S3. Rab32 is necessary to support Rab10 phosphorylation and Lamp1 levels**

**A)** Representative western data of siRNA-mediated knock down in B16 cells for Rab32-sub family of GTPases, Lrrk2, Lamp1 and pT73-Rab10/Rab10. Data is representative of three independent experiments.

**B)** Quantification of protein lysates analyzed in (**A**) Data presented as mean ±S.E.M. Statistical analysis was determined through a two-way ANOVA with a Tukey multiple comparisons post-hoc test, where significant findings are reported relative to NTC. **p* <0.05, ***p*<0.01, ****p*<0.001, *****p*<0.0001.

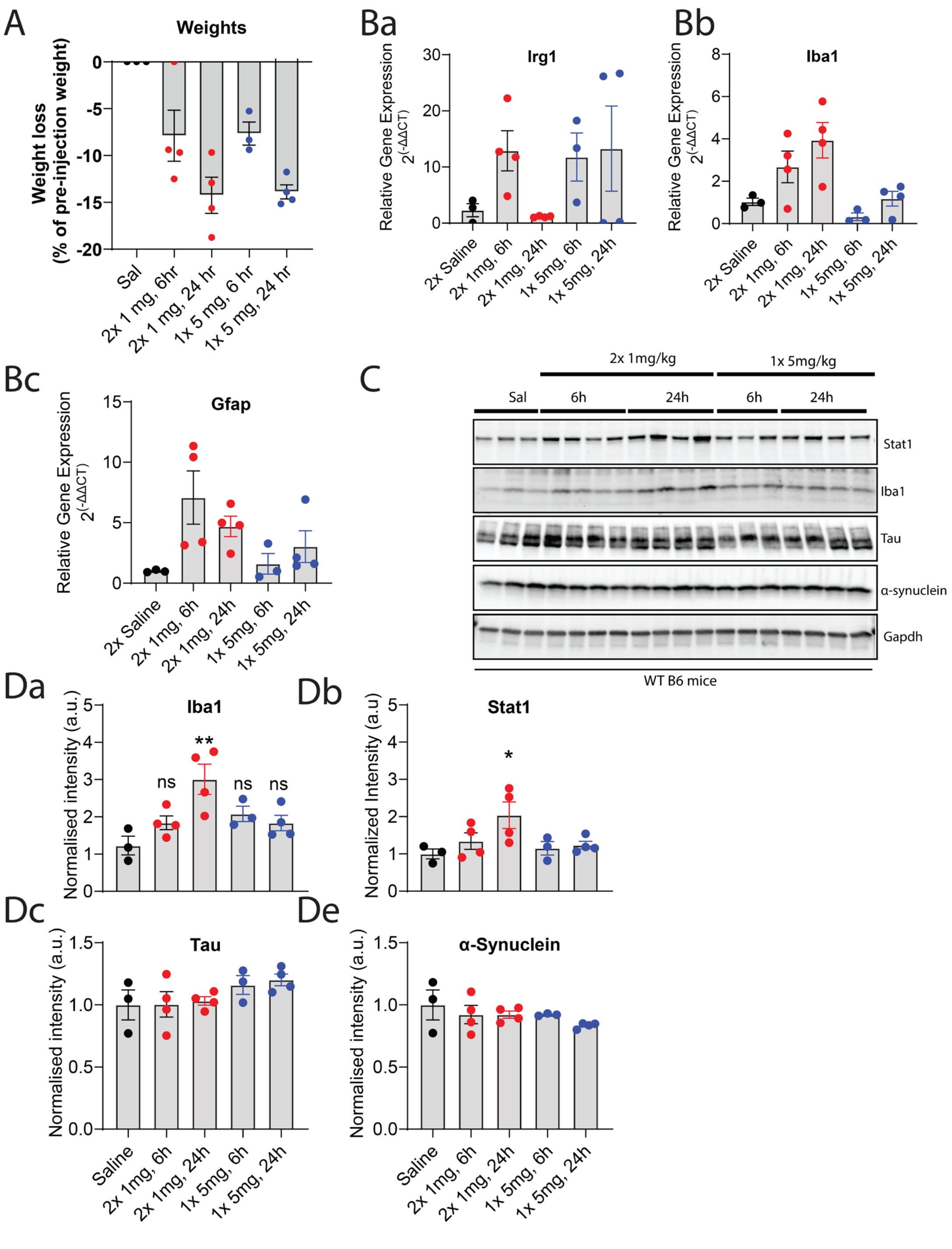


**Fig. S4. Establishing a working inflammatory paradigm in WT mice**

- **A)** Weight and weight loss (% of pre-injection weight) for three-month old WT mice that received either saline control, 1mg/kg or 5mg/kg LPS, for the dosing and time stated.
- **B)** Relative gene expression ($2^{(-\Delta\Delta CT)}$) of immunoresponsive genes from WT mice were assessed by qPCR after 6 and 24 hours post-injection with 1 or 5mg/kg of peripheral LPS or saline control. Data presented as mean ±S.E.M with eat dot representing an individual animal that underwent the treatment paradigm.

**C)** Western blot data of RIPA-soluble midbrain lysates from three-month old WT mice that received either saline control, 1mg/kg or 5mg/kg LPS.

**D)** Quantification of WB data presented in **C**. Each dot on the graph represents an individual animal that underwent the treatment paradigm. Data presented as mean ±S.E.M and analyzed by one-way ANOVA. *$p$ <0.05, **$p$<0.01, ***$p$<0.001, ****$p$<0.0001.

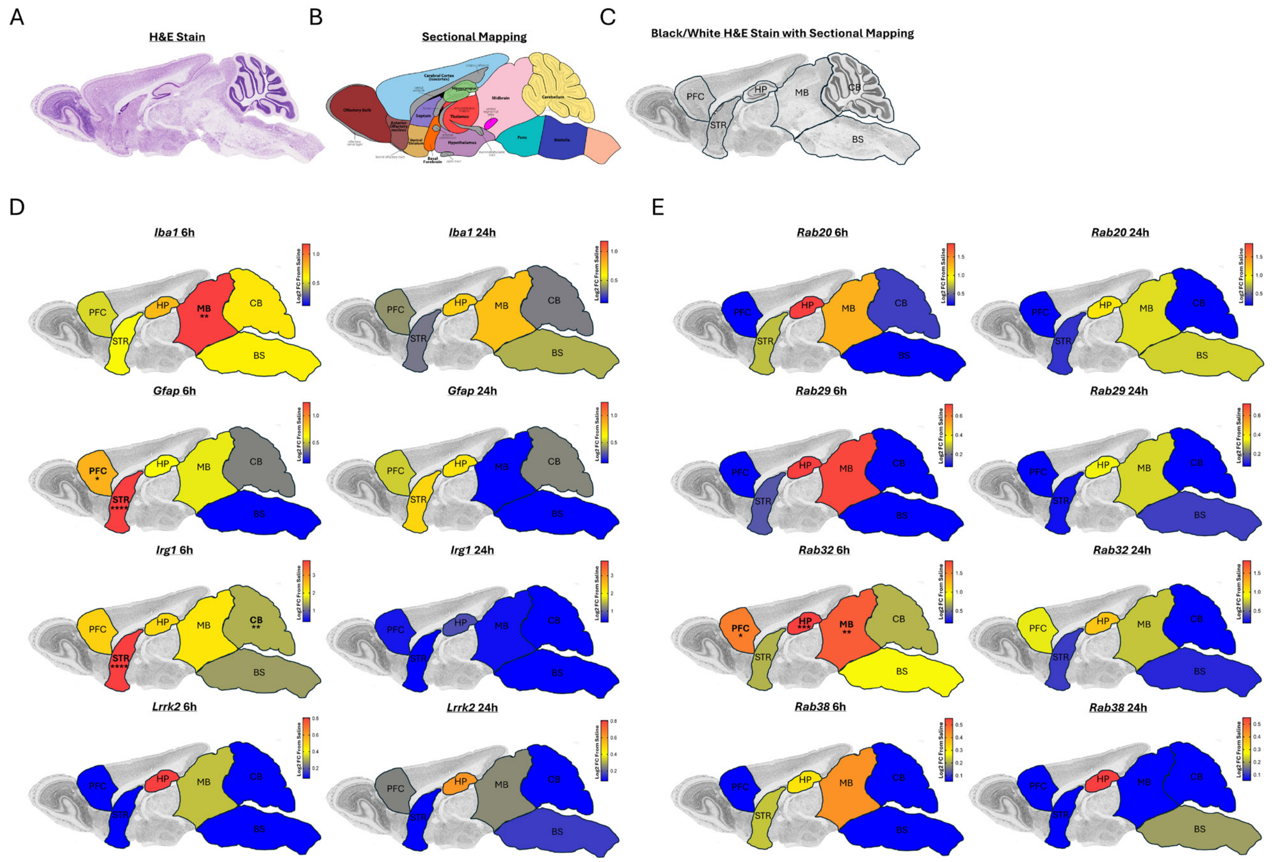


**Fig. S5. Sagittal ideogram of regional gene expression ($2^{(-\Delta\Delta CT)}$) of immunoresponsive and Rab GTPase genes in the brain of wild type mice with LPS**

- **A)** H&E stain used for **Figure 3** gene expression brain mapping.
- **B)** Sectional mapping used as example for **Fig. 3**.
- **C)** Black and white image of H&E stain with relevant sections drawn out for **Fig. 3**. A-C images were derived from the GENESTAT Project at The Rockefeller University.
- **D)** Relative gene expression ($2^{(-\Delta\Delta CT)}$) of immunoresponsive genes from the prefrontal cortex (PFC), striatum (STR), hippocampus (HP), midbrain (MB), cerebellum (CB) and brain stem (BS) of WT mice were assessed by qPCR after 6 and 24 hours post-injection with 1 mg/kg of peripheral LPS or saline control. Brain mapping to a sagittal ideogram of the mouse brain was created by determining Log2 fold change (FC) of $2^{(-\Delta\Delta CT)}$ of each examined gene between saline control and either 6 or 24 hour cohorts. Heatmaps of Log2 FC were created and transposed onto a sagittal mouse brain ideogram of an H&E stain (black and white colored) derived from the GENESTAT Project at The Rockefeller University (see **A-C** for original brain slice images used). Statistical analysis was determined through a two-way ANOVA with a Tukey multiple comparisons post-hoc test, where significant findings between saline control and 6 or 24 hour cohorts were reported by stars drawn on the brain region. *$p$ <0.05, **$p$<0.01, ***$p$<0.001, ****$p$<0.0001. Comparative statistical analysis of gene expression with/without peripheral LPS treatment (6 and 24 hour), within all 6 brain regions, is provided by two-way ANOVA (**Table S2**). The relative gene expression of each individual gene examined within each brain region by one-way ANOVA (**Fig. S6**).

**E)** Relative gene expression ($2^{(-\Delta\Delta CT)}$) of Rab GTPase genes from the prefrontal cortex (PFC), striatum (STR), hippocampus (HP), midbrain (MB), cerebellum (CB) and brain stem (BS) of WT mice were assessed by qPCR after 6 and 24 hours post-injection with 1 mg/kg of peripheral LPS or saline control. Brain mapping to a sagittal ideogram of the mouse brain was created by determining Log2 fold change (FC) of $2^{(-\Delta\Delta CT)}$ of each examined gene between saline control and either 6 or 24 hour cohorts. Heatmaps of Log2 FC were created and transposed onto a sagittal mouse brain ideogram derived from the GENESTAT Project (see **A-C** for original brain slice images used). Statistical analysis was determined through a two-way ANOVA with a Tukey multiple comparisons post-hoc test, where significant findings between saline control and 6 or 24 hour cohorts were reported by stars drawn on the brain region. *$p$ <0.05, **$p$<0.01, ***$p$<0.001, ****$p$<0.0001.. Comparative statistical analysis of gene expression with/without peripheral LPS treatment (6hr and 24hr), within all 6 brain regions, is provided by two-way ANOVA (**Table S2)**. The relative gene expression of each individual gene examined within each brain region by one-way ANOVA (**Fig. S6)**.

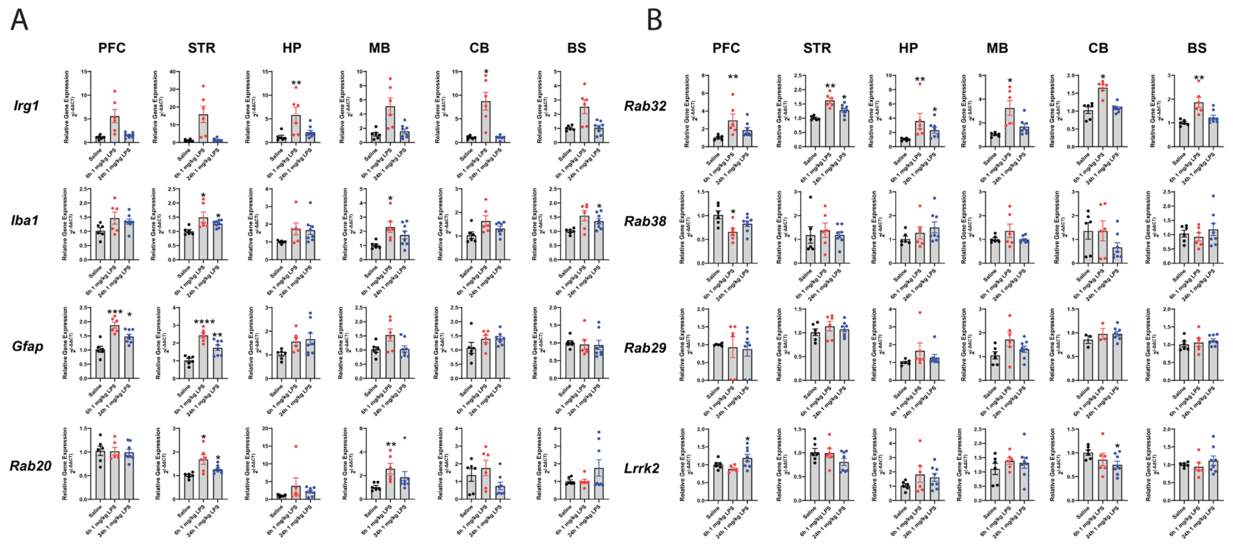


**Fig. S6. One-way ANOVA bar graphs of gene expression examined within each section of the mouse brain**

**A)** Relative gene expression ($2^{(-\Delta\Delta CT)}$) of immunoresponsive genes from the prefrontal cortex (PFC), striatum (STR), hippocampus (HP), midbrain (MB), cerebellum (CB) and brain stem (BS) of WT mice were assessed by qPCR after 6 and 24 hours post-injection with 1 mg/kg of peripheral LPS or saline control. Data presented as mean ±S.E.M and analyzed by one-way ANOVA (parametric: Brown-Forsythe and Welch ANOVA tests with Dunnett T3 multiple comparison's post-hoc test; non-parametric: Kruskal-Wallis test with a Dunn's multiple comparison post-hoc test).

**B)** Relative gene expression ($2^{(-\Delta\Delta CT)}$) of Rab GTPase genes from the prefrontal cortex (PFC), striatum (STR), hippocampus (HP), midbrain (MB), cerebellum (CB) and brain stem (BS) of WT mice were assessed by qPCR after 6 and 24 hours post-injection with 1 mg/kg of peripheral LPS or saline control. Data presented as mean ±S.E.M and analyzed by one-way ANOVA (parametric: Brown-Forsythe and Welch ANOVA tests with Dunnett T3 multiple comparison post-hoc test; non-parametric: Kruskal-Wallis test with a Dunn's multiple comparison post-hoc test).

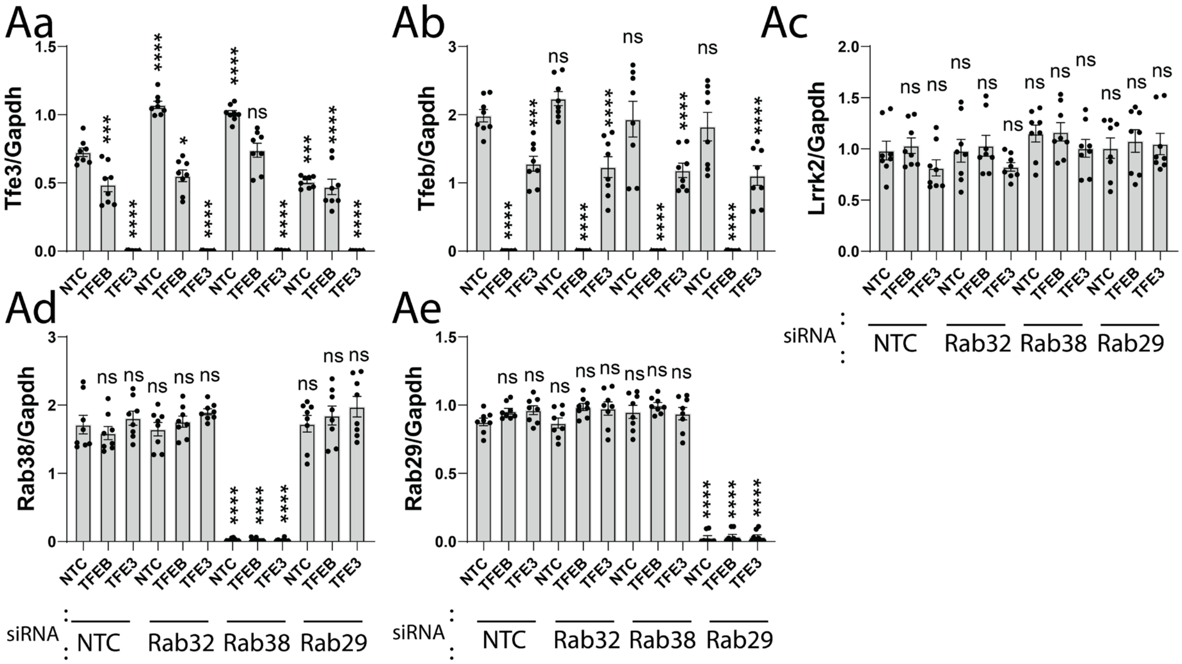


**Fig. S7. Bar graphs of protein markers related to Fig. 7**

**A)** Quantification of Tfe3, Tfeb, Lrrk2, Rab38 and Rab29 (**Aa-Ae**) normalized to Gapdh loading control, as presented in Figure 7. Data presented as mean ± S.E.M. Statistical analysis was determined through a two-way ANOVA with a Tukey multiple comparisons post-hoc test, where significant findings are reported as $*p<0.05$, $**p<0.01$, $***p<0.001$, $****p<0.0001$.

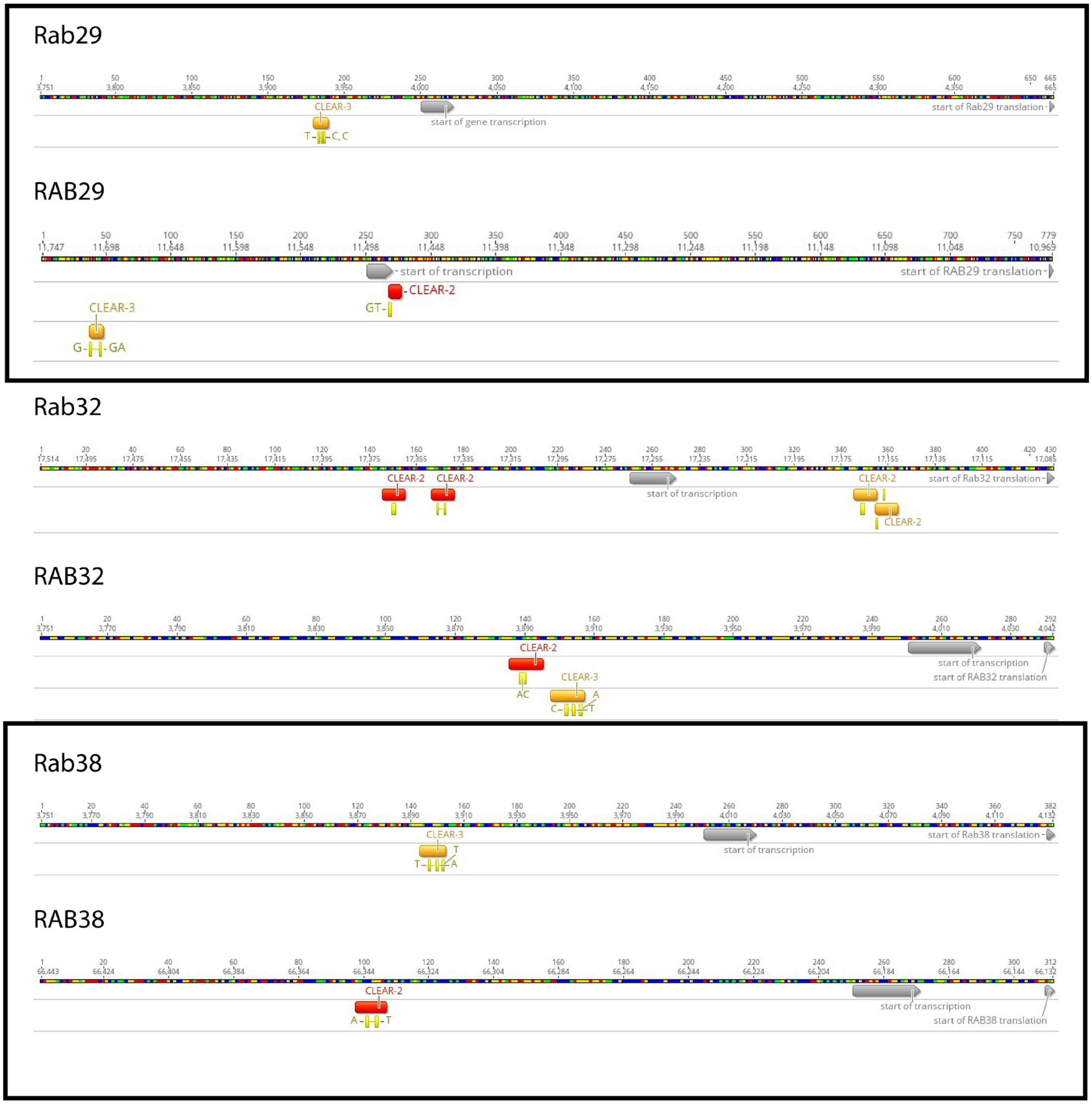


**Fig. S8 Comparative analysis of RAB29, 32 and 38 promoter sequences**

Promoter regions ~250bp upstream of the transcriptional start site (gray bar) to the start of translation (gray arrowhead) for murine and human RAB29, 32 and 38 genes. Sequences approximating the 10bp palindromic 'Coordinated Lysosomal Expression and Regulation' (CLEAR) consensus are highlighted in red (2bp mismatch) or orange (3bp mismatch), with 80% or 70% identity, respectively. The greatest affinity for transcription factor binding and gene expression is mediated by "tandem" CLEAR motifs within −195 to −118 base pairs 5' of the TSS *(1)* and only the promoters of Rab32/RAB32 approximate this configuration. Notably, Rab32, RAB32 have 5' CLEAR-2 sites that match criteria for E-box sequences (CANNTG) known to bind transcription factor 3 (TFE3)*(2)*.

Genomic sequences and annotations were obtained from the Ensembl database for murine C57BL/6J [GRCm39] Rab32 (Genomic ENSMUSG00000019832; transcript ENSMUST00000019974.5; Rab32-201), Rab38 (Genomic ENSMUSG00000030559; transcript ENSMUST00000107256; Rab38-201) and Rab29 (Genomic ENSMUSG00000026433; transcript ENSMUST00000027693; Rab29-201) and human [GRCh38.p14] RAB32 (Genomic ENSG00000118508; transcript ENST00000367495; RAB32-201), RAB38 (Genomic ENSG00000123892; transcript ENST00000243662; RAB38-201) and RAB29 (Genomic ENSG00000117280; transcript ENST00000367139; RAB29-201) loci (reference sequence identifiers).

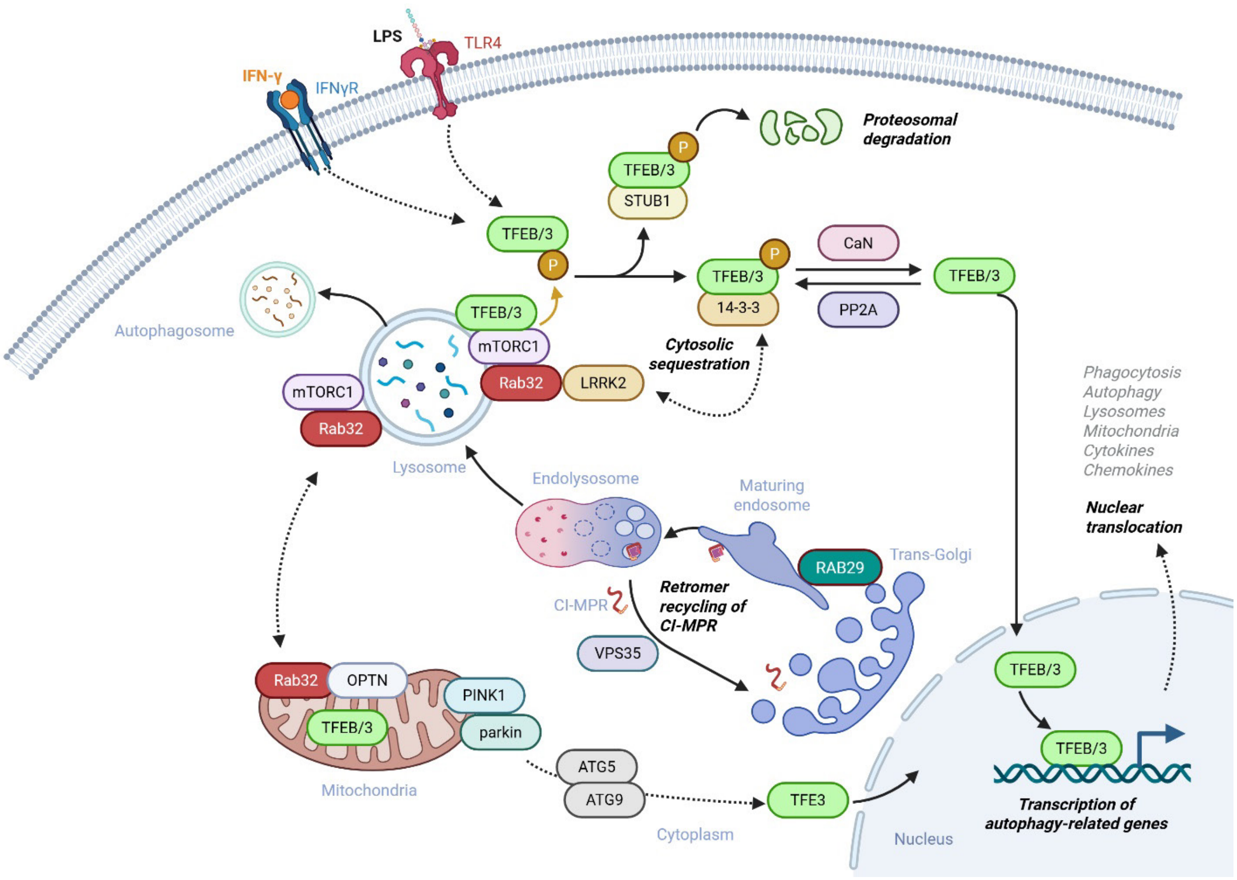


**Fig. S9: Graphical figure and abstract**

Ideogram of RAB32 in myeloid cells, its complex relationship with mTOR on lysosomes, MiT transcription factors in lysosomal biogenesis and mitophagy, focused on the role of Mendelian genes implicated in Parkinson's disease. In a replete state RAB32, in a GTP/GDP independent manner, is recruited by heterodimeric Rag GTPase to lysosomes. This induces mTORC1 kinase activity to phosphorylate MiT transcription factors, S321 TFE3 and S211 TFEB*(3)* that enables 14-3-3 binding and cytoplasmic sequestration and/or proteosomal degradation through interaction with STUB1*(4)*. LRRK2 kept dormant in cytoplasm as part of a 14-3-3 complex*(5)* is recruited to lysosomes in response to stress, inflammation or damage to lysosomal membranes *(6)*. The abundance of lysosomal LRRK2 and its kinase activity negatively regulate the nuclear localization of MiT transcription factors *(7)*. Conversely, lysosomal calcium signaling regulates autophagy through TFEB dephosphorylation by calcineurin *(8, 9)*. PP2A activation in response to oxidative stress may also trigger TFEB/3 dephosphorylation, inducing its nuclear translocation and subsequent transcriptional activity *(10, 11)*. Furthermore, RAB32 determines the targeting of PKA to mitochondria, increasing the phosphorylation of Drp1, regulating fission and mitophagy *(12–16)*. Concomitantly, Rab32 regulates Golgi structure and cell migration through PKA-mediated phosphorylation of OPTN *(17)*. In macrophages, LPS and IFNγ signaling activates TFEB, the mitochondrial TCA-cycle, itaconate synthesis *(18)* and delivery to nascent autophagosomes, tethered by RAB32 and LRRK2 *(19)*. Enhanced by mTOR inhibition, TFEB translocation to mitochondria then regulates complex 1 to down-modulate inflammation and the expression of pro-inflammatory cytokines *(20)*. MiT transcription factors are also activated during mitophagy downstream of PINK1 and Parkin *(21)*. Lastly, VPS35 is illustrated to highlight retromer's role in cation-independent receptor recycling. This is a prerequisite to enable the traffic of newly synthesized hydrolases from the trans-Golgi via maturing endosome to endolysosome and lysosome compartments *(22)*. RAB29 activates LRRK2 on Golgi and endosomal membranes *(23, 24)*, although its expression is not essential on lysosomes*(25)* and is tied to the expression of its homolog, RAB32, by MiT transcription.

### References

1. E. Brunialti, N. Rizzi, R. Pinto-Costa, A. Villa, A. Panzeri, C. Meda, M. Rebecchi, D. A. Di Monte, P. Ciana, Design and validation of a reporter mouse to study the dynamic regulation of TFEB and TFE3 activity through in vivo imaging techniques. *Autophagy* **20**, 1879–1894 (2024).

2. I. Aksan, C. R. Goding, Targeting the Microphthalmia Basic Helix-Loop-Helix–Leucine Zipper Transcription Factor to a Subset of E-Box Elements In Vitro and In Vivo. *Molecular and Cellular Biology* **18**, 6930–6938 (1998).

3. K. Drizyte-Miller, J. Chen, H. Cao, M. B. Schott, M. A. McNiven, The small GTPase Rab32 resides on lysosomes to regulate mTORC1 signaling. *Journal of Cell Science* **133** (2020), doi:10.1242/JCS.236661.

4. Y. Sha, L. Rao, C. Settembre, A. Ballabio, N. T. Eissa, STUB1 regulates TFEB-induced autophagy-lysosome pathway. *EMBO J* **36**, 2544–2552 (2017).

5. J. A. Martinez Fiesco, A. Beilina, A. Alvarez de la Cruz, N. Li, R. D. Metcalfe, M. R. Cookson, P. Zhang, 14-3-3 binding maintains the Parkinson's associated kinase LRRK2 in an inactive state. *Nat Commun* **16**, 7226 (2025).

6. A. Bentley-DeSousa, A. Roczniak-Ferguson, S. M. Ferguson, A STING–CASM–GABARAP pathway activates LRRK2 at lysosomes. *Journal of Cell Biology* **224**, e202310150 (2025).

7. N. Yadavalli, S. M. Ferguson, LRRK2 suppresses lysosome degradative activity in macrophages and microglia through MiT-TFE transcription factor inhibition. *Proceedings of the National Academy of Sciences of the United States of America* **120** (2023), doi:10.1073/PNAS.2303789120.

8. D. L. Medina, S. Di Paola, I. Peluso, A. Armani, D. De Stefani, R. Venditti, S. Montefusco, A. Scotto-Rosato, C. Prezioso, A. Forrester, C. Settembre, W. Wang, Q. Gao, H. Xu, M. Sandri, R. Rizzuto, M. A. De Matteis, A. Ballabio, Lysosomal calcium signalling regulates autophagy through calcineurin and TFEB. *Nat Cell Biol* **17**, 288–299 (2015).

9. S. Nakamura, S. Shigeyama, S. Minami, T. Shima, S. Akayama, T. Matsuda, A. Esposito, G. Napolitano, A. Kuma, T. Namba-Hamano, J. Nakamura, K. Yamamoto, M. Sasai, A. Tokumura, M. Miyamoto, Y. Oe, T. Fujita, S. Terawaki, A. Takahashi, M. Hamasaki, M. Yamamoto, Y. Okada, M. Komatsu, T. Nagai, Y. Takabatake, H. Xu, Y. Isaka, A. Ballabio, T. Yoshimori, LC3 lipidation is essential for TFEB activation during the lysosomal damage response to kidney injury. *Nat Cell Biol* **22**, 1252–1263 (2020).

10. J. A. Martina, H. I. Diab, O. A. Brady, R. Puertollano, TFEB and TFE3 are novel components of the integrated stress response. *EMBO J* **35**, 479–495 (2016).

11. J.-F. Zhao, N. Shpiro, G. Sathe, A. Brewer, T. J. Macartney, N. T. Wood, F. Negoita, K. Sakamoto, G. P. Sapkota, Targeted dephosphorylation of TFEB promotes its nuclear translocation. *iScience* **27**, 110432 (2024).

12. N. M. Alto, J. Soderling, J. D. Scott, Rab32 is an A-kinase anchoring protein and participates in mitochondrial dynamics. *Journal of Cell Biology* **158**, 659–668 (2002).

13. M. Bui, S. Y. Gilady, R. E. B. Fitzsimmons, M. D. Benson, E. M. Lynes, K. Gesson, N. M. Alto, S. Strack, J. D. Scott, T. Simmen, Rab32 modulates apoptosis onset and mitochondria-associated membrane (MAM) properties. *J. Biol. Chem.* **285**, 31590–31602 (2010).

14. P. Chen, Y. Lu, B. He, T. Xie, C. Yan, T. Liu, S. Wu, Y. Yeh, Z. Li, W. Huang, X. Zhang, Rab32 promotes glioblastoma migration and invasion via regulation of ERK/Drp1-mediated mitochondrial fission. *Cell Death Dis* **14**, 198 (2023).

15. H.-L. Zhang, Q. Cui, X.-T. Yu, Y.-X. Hou, R.-J. Ma, P.-S. Lu, Y. Wang, S.-C. Sun, H.-H. Wang, Rab32-based vesicles coordinate mitochondria and actin for spindle migration and organelle rearrangement in oocyte meiosis. *Journal of Advanced Research* (2025), doi:10.1016/j.jare.2025.05.001.

16. S.-L. Lu, S. Chen, K. Noda, Y. Li, C.-Y. Tsai, H. Omori, Y. Kato, Z. Zhang, B. Chen, K. Tokuda, T. Zheng, M. Wakita, E. Hara, M. Fukuda, Y. Wada, E. Morita, N. Uzawa, S. Murakami, T. Noda, Evidence that mitochondria in macrophages are destroyed by microautophagy. *Nat Commun* **16**, 8123 (2025).

17. K. M. Johnson, M. G. Marley, K. Drizyte-Miller, J. Chen, H. Cao, N. Mostafa, M. B. Schott, M. A. McNiven, G. L. Razidlo, Rab32 regulates Golgi structure and cell migration through Protein Kinase A–mediated phosphorylation of Optineurin. *Proceedings of the National Academy of Sciences* **122**, e2502971122 (2025).

18. E.-M. Schuster, M. W. Epple, K. M. Glaser, M. Mihlan, K. Lucht, J. A. Zimmermann, A. Bremser, A. Polyzou, N. Obier, N. Cabezas-Wallscheid, E. Trompouki, A. Ballabio, J. Vogel, J. M. Buescher, A. J. Westermann, A. S. Rambold, TFEB induces mitochondrial itaconate synthesis to suppress bacterial growth in macrophages. *Nat Metab* **4**, 856–866 (2022).

19. H. Lian, D. Park, M. Chen, F. Schueder, M. Lara-Tejero, J. Liu, J. E. Galán, Parkinson's disease kinase LRRK2 coordinates a cell-intrinsic itaconate-dependent defence pathway against intracellular Salmonella. *Nature microbiology* **8**, 1880–1895 (2023).

20. C. Calabrese, H. Nolte, M. R. Pitman, R. Ganesan, P. Lampe, R. Laboy, R. Ripa, J. Fischer, R. Polara, S. K. Panda, S. Chipurupalli, S. Gutierrez, D. Thomas, S. M. Pitson, A. Antebi, N. Robinson, Mitochondrial translocation of TFEB regulates complex I and inflammation. *EMBO Rep* **25**, 704–724 (2024).

21. C. L. Nezich, C. Wang, A. I. Fogel, R. J. Youle, MiT/TFE transcription factors are activated during mitophagy downstream of Parkin and Atg5. *J Cell Biol* **210**, 435–450 (2015).

22. Y. Cui, Z. Yang, N. Flores-Rodriguez, J. Follett, N. Ariotti, A. A. Wall, R. G. Parton, R. D. Teasdale, Formation of retromer transport carriers is disrupted by the Parkinson disease-linked Vps35 D620N variant. *Traffic* **22**, 123–136 (2021).

23. E. Purlyte, H. S. Dhekne, A. R. Sarhan, R. Gomez, P. Lis, M. Wightman, T. N. Martinez, F. Tonelli, S. R. Pfeffer, D. R. Alessi, Rab29 activation of the Parkinson's disease-associated LRRK2 kinase. *The EMBO Journal* **37**, 1–18 (2018).

24. E. G. Vides, A. Adhikari, C. Y. Chiang, P. Lis, E. Purlyte, C. Limouse, J. L. Shumate, E. Spínola-Lasso, H. S. Dhekne, D. R. Alessi, S. R. Pfeffer, A feed-forward pathway drives LRRK2 kinase membrane recruitment and activation. *eLife* **11** (2022), doi:10.7554/ELIFE.79771.

25. A. F. Kalogeropulou, J. B. Freemantle, P. Lis, E. G. Vides, N. K. Polinski, D. R. Alessi, Endogenous Rab29 does not impact basal or stimulated LRRK2 pathway activity. *Biochemical Journal* **477**, 4397–4423 (2020).
